# Chromosome Pairing Through Tensed DNA Tethers Model Revealed by BRCA2 Meiotic Domain Deletion

**DOI:** 10.1101/2023.10.06.561239

**Authors:** Lieke Koornneef, Sari E. van Rossum-Fikkert, Esther Sleddens-Linkels, Simona Miron, Alex Maas, Yvette van Loon, Marco Barchi, Sophie Zinn-Justin, Jeroen Essers, Roland Kanaar, Willy M. Baarends, Alex N. Zelensky

**Affiliations:** Department of Developmental Biology, Erasmus University Medical Center, 3000 CA, Rotterdam, The Netherlands; Oncode Institute, Erasmus University Medical Center, 3000 CA, Rotterdam, The Netherlands; Department of Molecular Genetics, Erasmus MC Cancer Institute, Erasmus University Medical Center, 3000 CA, Rotterdam, The Netherlands; Institute for Integrative Biology of the Cell (I2BC), CEA, CNRS, Uni Paris-Sud, Uni Paris-Saclay, Gif-sur-Yvette, France; Department of Cell Biology, Erasmus University Medical Center, 3000 CA, Rotterdam, The Netherlands; Deparment of Biomedicine and Prevention, Faculty of Medicine, University of Rome Tor Vergata, Rome, Italy; Department of Radiotherapy, Erasmus University Medical Center, 3000 CA Rotterdam, The Netherlands; Department of Vascular Surgery, Erasmus University Medical Center, 3000 CA, Rotterdam, The Netherlands

## Abstract

BRCA2 has multiple functional domains that interact with different partners, and is essential for both somatic and meiotic homologous recombination (HR). We created a *Brca2*^*Δ12-14*^ mouse model with an internal deletion of the region which we named “the meiotic domain of BRCA2”, as its loss results in complete failure of meiotic HR, while somatic HR is intact. The deletion in the protein includes the HSF2BP-binding motifs (exons 12-13) and the DMC1-binding PhePP domain (exon 14). *Brca2*^*Δ12-14*^ mice showed complete infertility in both males and females, with sexually dimorphic features. Recombinase foci (both RAD51 and DMC1) were completely undetectable in mutant spermatocytes, but while DMC1 foci were also absent in mutant oocytes, RAD51 foci numbers were only partially reduced. The function of the PhePP domain for meiotic HR is unclear, but both the phenotype of *Brca2*^*Δ12-14*^, and our biochemical data indicate that, along with the BRC repeats of BRCA2, PhePP is both critical and specific for DMC1 loading in meiotic HR, analogous to the C-terminal RAD51-specific TR2/CTRB. Further investigation of DSB end processing in *Brca2*^*Δ12-14*^ meiocytes and controls, using super-resolution imaging of RPA and SYCP3 led to discovery of two novel features. First, in Brca2*Δ*12-14 oocytes, but not in the spermatocytes nor wild types, we observed RPA foci as doublets ∼200 nm apart, which could represent DSB end resolution into separate nanofoci. Second, we describe RPA structures that are completely HR-dependent and are indicative of long, double-stranded DNA connections between homologs prior to synapsis. Our observations lend support to a model for chromosome alignment via multiple HR-dependent DNA tethers that connect homologs and may be tensed. We propose that tether shortening (e.g. by dynamic adjustment of chromatin loops by meiotic cohesins) provides a plausible molecular mechanism to juxtapose homologs and initiate synapsis.

**Version 2 Revision Summary:** The main difference compared to version 1 (deposited on October 6 2023) is a more concise and structured description of the tensed DNA tether model of meiotic chromosome pairing, based on the discussions with colleagues and one round of peer review. In the new model presentation, we explicitly separated the inferences from the presented data from the two hypothetical propositions: (1) tether shortening contributes to pairing rather than simply accompanies it, and (2) the apparent tension, which reveals the tethers on chromosome spreads, also exists in the nuclei. We also clarified the definition of the tether, avoiding the ambiguous “RPA tether” term, and provided a more complete overview of the relevant prior literature on proteinaceous bridges and DNA connections. Biochemical data (Fig. 4, S5) has been replicated under uniform conditions and extended to mouse proteins. Manuscript has been reformatted to improve on-screen readability.

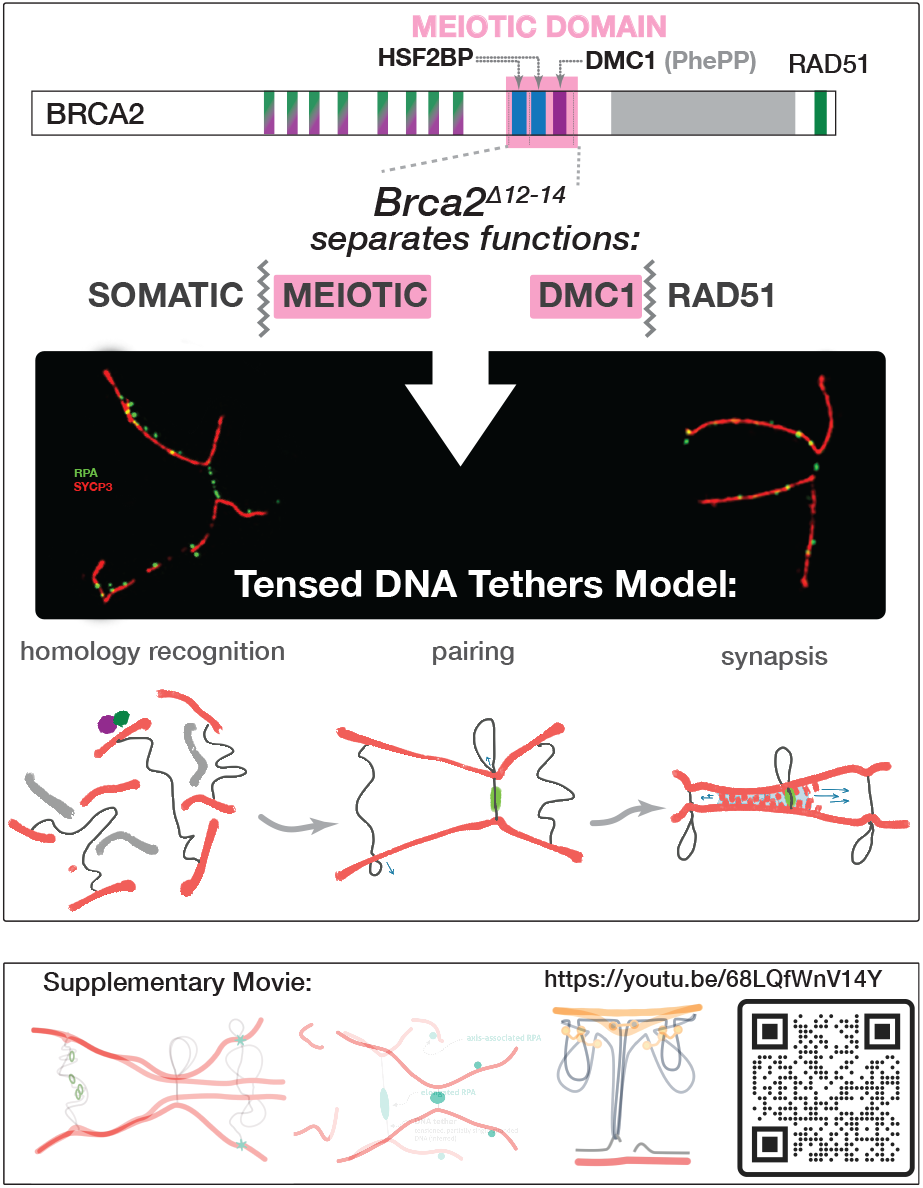

## INTRODUCTION

BRCA2 is essential for both mitotic and meiotic homologous recombination (HR) in vertebrates ^1,2^. Its meiotic role is evolutionarily ancestral: loss of fertility due to meiotic HR failure is the universal and the most pronounced phenotype resulting from the deletion of BRCA2 orthologues in fungi^3^, plants^4,5^, protozoa ^6^ and invertebrates ^7–10^. However, few details are available about the exact meiotic function of BRCA2. There is no BRCA2 orthologue in budding yeast, a key model system for mechanistic studies of meiosis. Also, in vertebrates, studies of BRCA2 in meiosis are limited by embryonic lethality of *Brca2* knockout in mouse and difficulty of immunodetection of the protein on meiotic chromosome spreads. Thus, the mechanistic understanding of BRCA2 in meiosis is mostly based on inferences from the studies of mitotic HR and limited biochemical experiments. Still, three mouse models confirmed the critical role of *Brca2* in mouse meiotic HR: a knockout lethality rescued by a hypomorphic transgene that is not active in meiocytes ^11^ and two mouse strains with different truncating mutations in exon 11 ^12,13^. In all cases complete infertility of both sexes was observed. While BRCA2-mediated loading of RAD51 in somatic cells is required for repair of accidental DNA damage, loading of RAD51 and its meiosis-specific paralog DMC1 in meiosis is essential for the repair of programmed DNA double-strand breaks (DSBs). This special form of HR repair is then pivotal to achieve complete and stable homologous chromosome pairing and synapsis (formation of the synaptonemal complex connecting the homologous chromosomes), as well as for crossover formation.

Studies on BRCA2 in somatic cells were focused on its functional interaction with RAD51, the recombinase that performs homology recognition and strand exchange during HR. BRCA2 also interacts with DMC1 ^14–17^, the meiosis-specific paralog of RAD51. The BRCA2 BRC repeats encoded by *BRCA2* exon 11 interact with both recombinases, and there are additional sites that interact with only one: the RAD51-binding site in the C-terminal domain of BRCA2 encoded by exon 27, and the DMC1-binding site located before the highly conserved DNA-binding domain and encoded by exon 14 ^16^. The data on this DMC1-specific interaction domain is contradictory: it was identified biochemically using recombinant proteins ^16^, but its physiological relevance was later put into question when a mouse strain with a nonconservative substitution of phenylalanine 2351 to aspartate in BRCA2 was engineered ^18^. The mouse F2351 corresponds to F2406 in human BRCA2, which was absolutely essential for the interaction between recombinant human BRCA2 and DMC1 *in vitro*. However, the *Brca2*^*F2351D*^ mice were fully fertile.

We and others have recently discovered that BRCA2 forms a high-affinity complex with the previously uncharacterized protein HSF2BP (also called MEILB2) in mouse embryonic stem (mES) cells and meiocytes ^19–24^. Mice deficient for HSF2BP are born at Mendelian ratios and have no overt somatic phenotypes, but males are infertile due to meiotic HR failure. Shortly after, HSF2BP was shown to directly interact with another uncharacterized protein named BRME1 (also called MAMERR or MEIOK21), and the phenotypes of the *Hsf2bp* and *Brme1* knockout mouse models ^19,21,25–29^ were similar.

To test whether HSF2BP interaction with BRCA2 is crucial for its meiotic function, we previously generated and characterized a mouse strain lacking *Brca2* exon 12 ^22^, which encodes one of the two motifs comprising the HSF2BP-binding domain (HBD, Figure 1A) of BRCA2, the other motif is localized in exon 13. As exon 12 boundaries fall in the same position relative to the reading frame, this deletion did not lead to a frame shift. The *Brca2*^*Δ12*^ mice were phenotypically normal and fertile, in contrast to the male sterility of the *Hsf2bp* knockout ^19,21,25^. This put the role of HSF2BP-BRCA2 interaction in meiotic HR into question. However, while exon 12 deletion reduced HSF2BP-BRCA2 interaction, it did not completely abolish it *in vivo*. The residual affinity was higher in the testis co-immunoprecipitation experiments than in biochemical experiments with purified or ectopically produced proteins, suggesting that additional factors may contribute to the HSF2BP-BRCA2 interaction in the physiological context. In the same work, we also characterized mES cells with a deletion of exons 12 through 14 of *Brca2* (*Δ*12-14). Exon 14 is the nearest downstream exon that can be co-excised with exon 12 while preserving the reading frame. In addition to the two HSF2BP-binding motifs encoded by exons 12 and 13, deletion of exon 14 removes a block of conserved amino acids that includes the site that has previously been shown to bind to DMC1 (the PhePP motif ^16^), as mentioned above. In these cells HSF2BP did not co-precipitate at all with the BRCA2 produced from the deletion allele.

**Figure 1.**
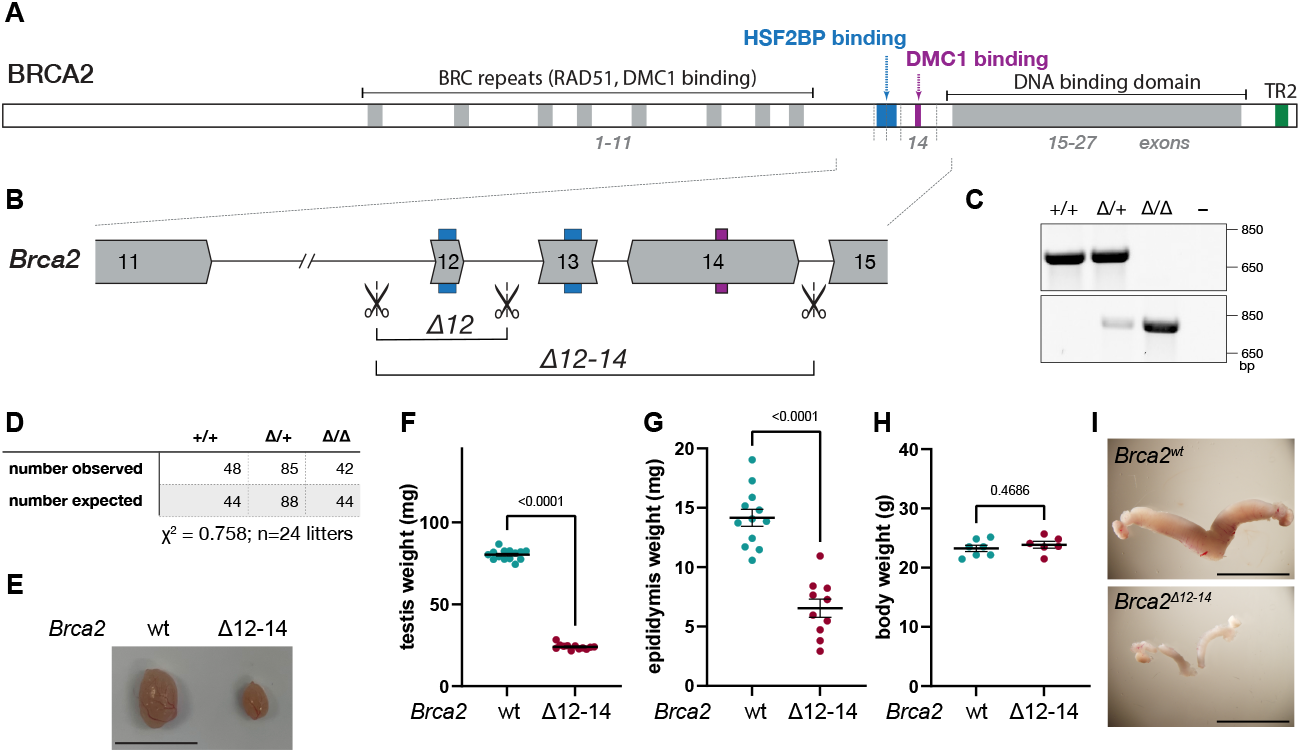
Characterization of *Brca2*^*Δ12-14*^ mouse model. **(A)** Schematic domain structure of BRCA2 protein. **(B)** Exon-intron structure of the region targeted by CRISPR/Cas9 deletion. **(C)** RT-PCR and immunoblot analysis confirming the effect of exon 12-14 deletion on *Brca2* transcript and BRCA2 protein. **(D)** Numbers of pups born from intercrosses between *Brca2*^*Δ12-14/*+^ animals and comparison to the numbers expected if Mendelian ratios are maintained. **(E)** Representative photographs of testes from *Brca2*^*wt*^ and *Brca2*^*Δ12-14*^ littermate animals. Testis **(F)**, epididymis **(G)** and body weight **(H)** in *Brca2*^*Δ12-14*^ and control male mice (n=6-7, n=6-7 and n=5-6 respectively). All data is plotted, mean, s.e.m. and *p* values from two-tailed unpaired t-test are indicated. **(I)** Representative photographs of ovaries and uterus from *Brca2*^*wt*^ and *Brca2*^*Δ12-14*^ littermate animals of 18 weeks old. Scale bar represents 1 cm **(E**,**I)**.

Here we describe the phenotype of the mouse model with *Brca2* exon 12-14 deletion, which lacks both HSF2BP-binding domains and the purported DMC1-binding site. The strain is viable but completely infertile, thus defining the deleted region as the “meiotic domain” of BRCA2. The phenotype provides novel insights into BRCA2 structure-function relationships in terms of somatic versus meiotic functions, and male versus female functions. Based on super-resolution microscopy analysis of meiotic chromosome spreads we discuss a model that postulates a role for DNA tethers in homologous chromosome pairing and synapsis and the potential role of tension in this.

## RESULTS

### No somatic defects and complete meiotic arrest in Brca2^Δ12-14^ mouse

To test whether the complete removal of the BRCA2 HSF2BP-binding domain recapitulates the *Hsf2bp* knockout phenotype, we engineered a homozygous mouse strain lacking *Brca2* exons 12-14 (*Brca2*^*Δ12-14*^), following the strategy we previously tested in mES cells ^22^ (Figure 1A-C). Consistent with the lack of HR defects in mES cells carrying the same mutation (Figure S1 and ref ^22^), *Brca2*^*Δ12-14*^ mice were viable, of normal appearance, and were born at Mendelian ratios (Figure 1D). However, *Brca2*^*Δ12-14*^ males as well as *Brca2*^*Δ12-14*^ females were infertile, demonstrated by the lack of successful breedings after mating 8 *Brca2*^*Δ12-14*^ males and 6 *Brca2*^*Δ12-14*^ females with their corresponding wild type partners. In males, *Brca2*^*Δ12-14*^ testes were smaller (Figure 1E) with 70% reduced weight compared to wild type (Figure 1F). Epididymis weight was also reduced by 54%, whereas body weight was unaffected (Figure 1G-H). Histological analyses of *Brca2*^*Δ12-14*^ testes showed that development of germ cells did not proceed beyond the spermatocyte stage (Figure S2A). In adult females we observed a reduction in size of the internal reproductive organs (Figure 1I). Histology of adult ovaries showed absence of follicles (Figure S2B). Together, this shows that exons 12-14 of *Brca2* are essential for fertility in both male and female mice.

### Synapsis in Brca2^Δ12-14^ spermatocytes is aberrant but more extensive compared to that of Spo11 knockout

Next, we studied the cause of *Brca2*^*Δ12-14*^ mice infertility using immunofluorescent staining for meiotic prophase proteins on *Brca2*^*Δ12-14*^ meiocyte spread nuclei and corresponding wild type littermate controls, which revealed major defects. Starting with staining for axial/lateral filaments of the synaptonemal complex (SYCP2 or SYCP3), we observed that *Brca2*^*Δ12-14*^ spermatocytes failed to perform faithful synapsis, resulting in a late zygotene-like appearance of the most advanced cell types (Figure S3A,B). In regions where synapsis occurred, SYCP1 (which forms the transverse filaments of the SC) was present, as expected (Figure S3C). Similar observations were made in E17.5 *Brca2*^*Δ12-14*^ oocytes (Figure S3D,E).

To investigate synapsis progression in spermatocytes in more detail, we used structured illumination microscopy (SIM) to study SYCP3. First, we observed that 91% of the partially synapsed regions in the mutant included telomeric ends, while this was only 66% for wild type, where partial synapsis was more often observed only in a central region, excluding the telomeres (p < 0.001, unpaired two-sided t-test, fully synapsed chromosomes were excluded) (Figure S3F,G). Second, in addition to the extensive aberrant multi-chromosome synapsis, apparently completely synapsed configurations were also clearly detected (Figure S3H), with a frequency that was two times higher in our *Brca2*^*Δ12-14*^ model than in a model without DSB formation (*Spo11*^−*/* −^ mutant) (Figure S3I). We used fluorescent in situ hybridization (FISH) of chromosome 13 in combination with immunocytochemistry of SYCP3 to assess whether this complete end-to-end synapsis was homologous. In only 7.4% of the analysed nuclei we observed a single FISH cloud that located on a fully synapsed chromosome (Figure S3J, K) while for or all other nuclei another FISH cloud in a different region was detected, or the single FISH cloud located to regions without synapsis or regions with unequal and partial synapsis This shows that in absence of exon 12-14 of *Brca2*, homologous chromosome synapsis is abrogated, and, surprisingly, that non-homologous synapsis occurs more extensively than in the absence of DSB formation.

### Brca2^Δ12-1^4 differentially affects DMC1 and RAD51 in oocytes and spermatocytes

As homologous chromosome synapsis is dependent on HR ^11,30–33^, and BRCA2 acts as HR mediator, we next analyzed the localization of RAD51 and DMC1 recombinases in spermatocyte and oocyte nuclei. In *Brca2*^*Δ12-14*^ spermatocytes, both RAD51 and DMC1 foci were completely absent (Figure 2A-C). To exclude defects in DSB formation, we assessed loading of the single-stranded DNA binding protein RPA as marker for DSB formation. In the mutant, RPA foci were present and the number was increased compared to wild type (Figure 2D,E), suggesting that DSBs can be formed, but that RPA cannot be replaced by recombinases. Also, the foci numbers of SPATA22, a meiosis-specific single-stranded DNA binding protein ^34– 36^, were higher in absence of exons 12-14 of *Brca2* than in the controls (Figure 2D,F), similar to what was observed in *Hsf2bp*^−*/* −^ and *Brme1*^−*/* −^ spermatocytes ^19,25,27,28^. Since the *Brca2*^*Δ12-14*^ mouse model completely lacks the HSF2BP-binding domain, we investigated the localization patterns of HSF2BP and its interaction partner BRME1. Both proteins were present in foci but behaved differently in the mutant (Figure 2G). HSF2BP foci numbers were reduced, while BRME1 foci numbers were increased (Figure 2H,I). Moreover, the intensity of the RPA, BRME1 and SPATA22 foci was increased (Figure S4A), while the intensity of HSF2BP was unaffected in *Brca2*^*Δ12-14*^ spermatocytes (Figure S4B).

**Figure 2.**
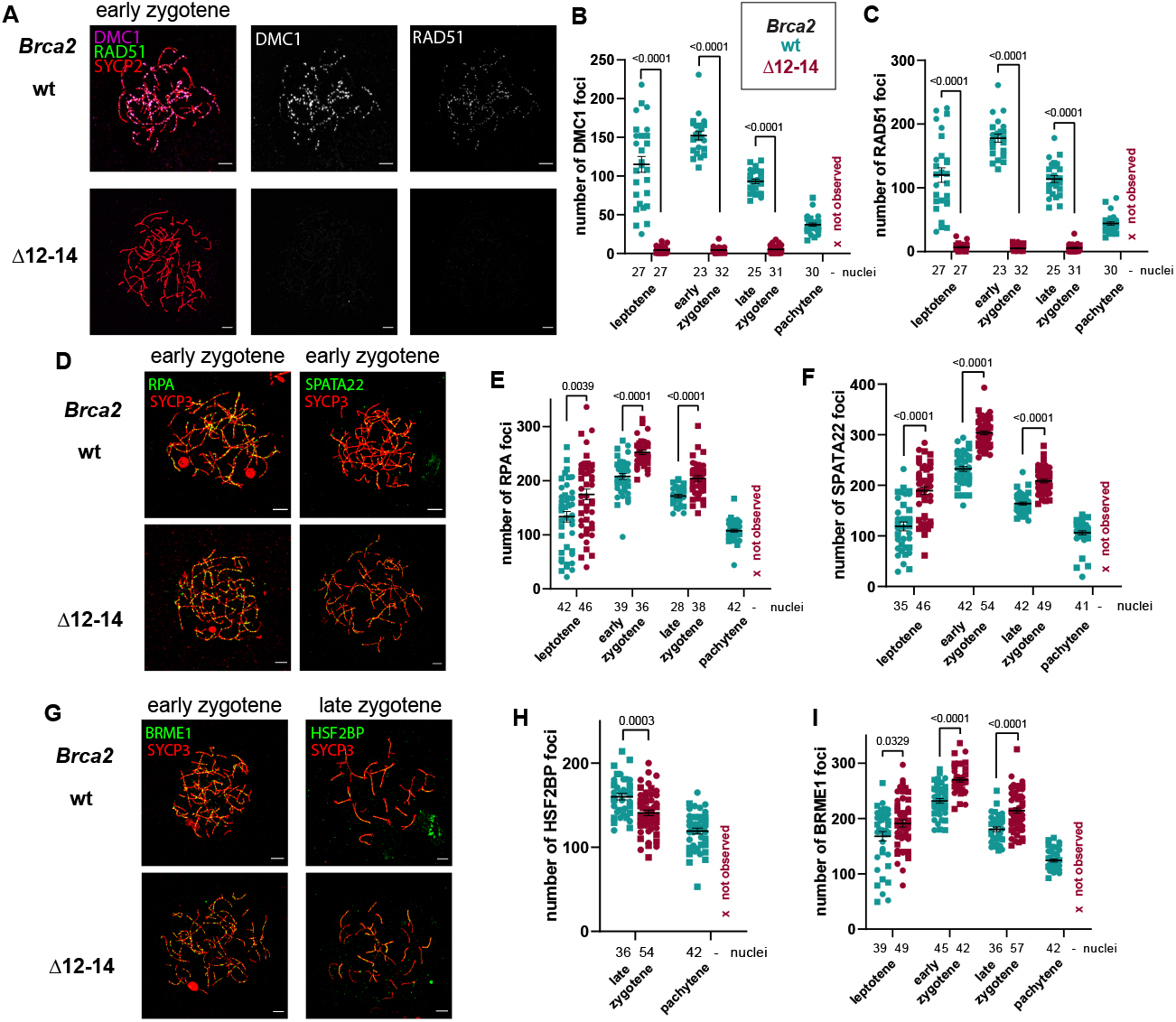
Meiotic analysis of *Brca2*^*Δ12-14*^ spermatocytes. Immunofluorescent analysis of meiotic proteins on nuclear surface spread spermatocytes from *Brca2*^*Δ12-14*^ and control mice. **(A)** Representative images of spread spermatocyte nuclei immunostained for RAD51 (green), DMC1 (magenta), and SYCP2 (red) including individual presentation of RAD51 or DMC1 in white. **(B)** Quantification of DMC1 foci. **(C)** Quantification of RAD51 foci. **(D)** Representative images of spread spermatocyte nuclei immunostained for RPA (green) or SPATA22 (green), and SYCP3 (red). **(E)** Quantification of RPA foci. **(F)** Quantification of SPATA22 foci. **(G)** Representative images of spread spermatocyte nuclei immunostained for BRME1 (green) or HSF2BP (green), and SYCP3 (red). **(H)** Quantification of HSF2BP foci. **(I)** Quantification of BRME1 foci. Mean, s.e.m., p-values from two-tailed unpaired t-test, and number of analyzed nuclei are indicated in the graphs. Symbol shapes represents individual animals: n = 2 for *Brca2*^*wt*^ and n= 2 for *Brca2*^*Δ12-14*^. Sea blue color represents *Brca2*^+*/*+^ and burgundy color represents *Brca2*^*Δ12-14*^. For the mutant the pachytene stage was not observed. Scale bar represents 5 µm **(A**,**D**,**G)**.

*Hsf2bp*^−*/* −^ and *Brme1*^−*/* −^ mice showed a sexual dimorphism regarding defects in meiosis ^19,21,25–29,37^. Therefore, we also analyzed meiotic HR protein localization patterns in embryonic (E15.5 and E17.5) *Brca2*^*Δ12-14*^ oocyte nuclei, since female meiotic prophase initiates during embryogenesis. Similar to the male mutant, DMC1 foci were not observed (Figure 3A,B). However, RAD51 foci were detected (Figure 3A), but their number in leptotene-like nuclei was 60% lower than in wild type leptotene (Figure 3C) and the foci were 20% less intense (Figure S4C). RAD51 foci numbers remained constant between early zygotene and late zygotene (Figure S4D), which distinguishes this mutant from *Hsf2bp*^*-/-*^ oocytes where, despite similar initial RAD51 foci numbers as in *Brca2*^*Δ12-14*^ oocytes, RAD51 foci numbers decreased as cells progressed during meiotic prophase ^25^ (Figure S4D). This could imply that meiotic HR repair progresses to a point where RAD51 protein is removed in *Hsf2bp*^−*/* −^ oocytes, but not in *Brca2*^*Δ12-14*^ oocytes. In addition, and similar and consistent with *Brca2*^*Δ12-14*^ males, RPA foci were present in *Brca2*^*Δ12-14*^ oocytes and numbers were increased (Figure 3D,E), confirming formation of DSBs. Also, SPATA22 foci numbers were increased in absence of exons 12-14 of *Brca2* (Figure 3D,F). Like SPATA22, HSF2BP and BRME1 showed similar changes as observed in the mutant spermatocytes: reduced HSF2BP, and increased BRME1 foci numbers (Figure 3G-I). Furthermore, the HSF2BP intensity was reduced, while the RPA, BRME1 and SPATA22 foci intensity was increased in the mutant oocytes (Figure S4E,F).

**Figure 3.**
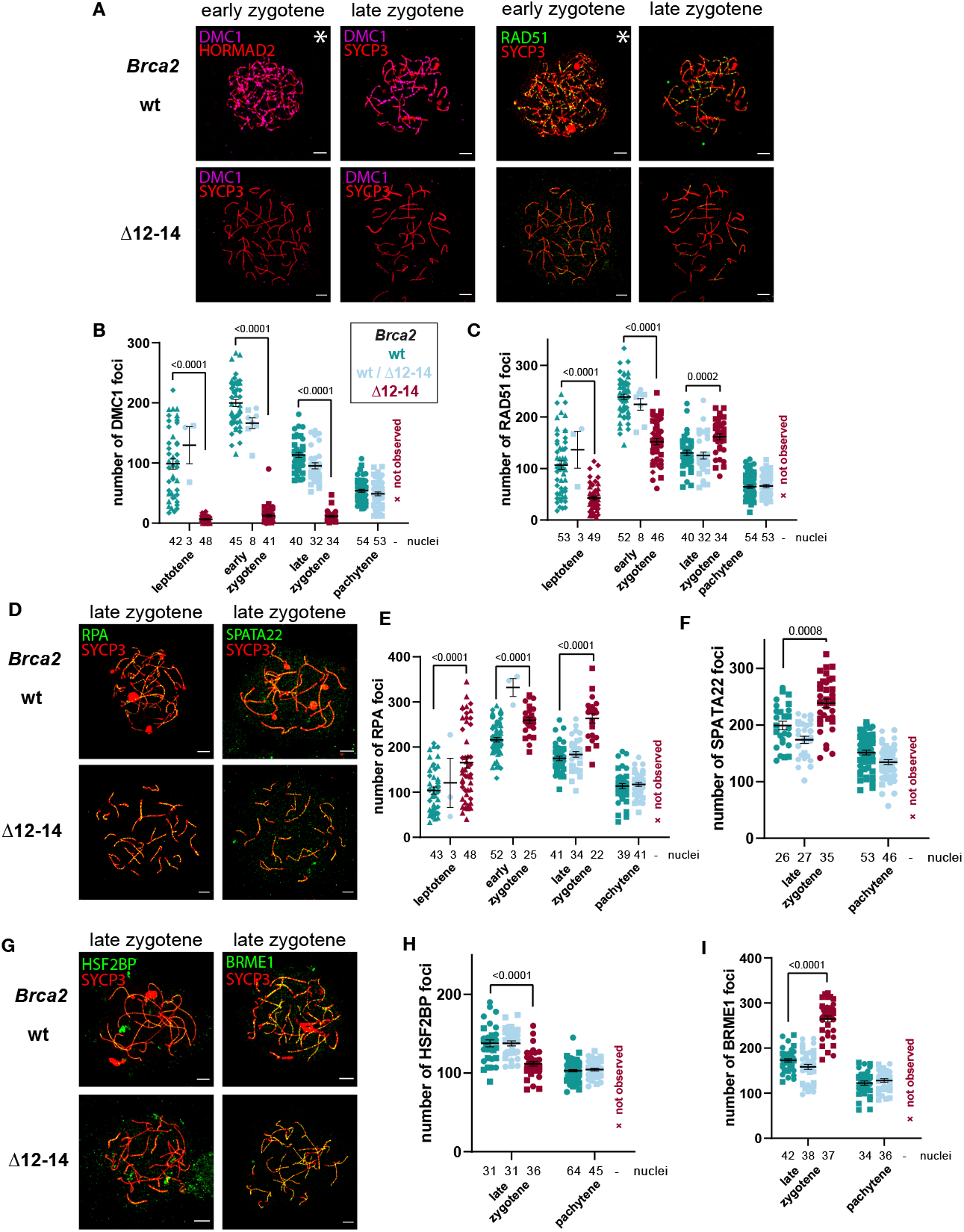
Meiotic analysis of *Brca2*^*Δ12-14*^ oocytes. Immunofluorescent analysis of meiotic proteins on nuclear surface spread oocytes from E15.5 and E17.5 *Brca2*^*Δ12-14*^ and control mice. **(A)** Representative images of spread spermatocyte nuclei immunostained for RAD51 (green) and DMC1 (magenta) with SYCP3 or HORMAD2 (red). Meiotic stage of nuclei is indicated above the images. For wild type late zygotene and pachytene and *Brca2*^*Δ12-14*^ early zygotene and late zygotene, oocytes spreads of E17.5 mice were used. For the earlier stages (leptotene and early zygotene of wild type and leptotene of *Brca2*^*Δ12-14*^) oocytes spreads of E15.5 mice were used. Images from E15.5 spreads are indicated with asterisks. **(B)** Quantification of DMC1 foci. **(C)** Quantification of RAD51 foci. **(D)** Representative images of spread oocyte nuclei immunostained for RPA (green) or SPATA22 (green), and SYCP3 (red). **(E)** Quantification of RPA foci. **(F)** Quantification of SPATA22 foci. **(G)** Representative images of spread oocyte nuclei immunostained for BRME1 (green) or HSF2BP (green), and SYCP3 (red). **(H)** Quantification of HSF2BP foci. **(I)** Quantification of BRME1 foci. Mean, s.e.m., p-values from two-tailed unpaired t-test, and number of analyzed nuclei are indicated in the graphs. Symbol shapes (circle, square, hexagon) represent individual E17.5 animals: n = 2 for *Brca2*^*wt*^, n = 2 for *Brca2*^*wt/Δ12-14*^ (n = 3 for SPATA22) and n = 2 for *Brca2*^*Δ12-14*^. Data and images of E15.5 oocytes originate from 2 animals of *Brca2*^*wt*^ and 2 animals of *Brca2*^*Δ12-14*^, which are represented in the graphs by the symbols rhombus and triangle. Sea blue color represents *Brca2*^*wt*^, light blue color represents *Brca2*^*wt/Δ12-14*^, burgundy color represents *Brca2*^*Δ12-14*^. For the mutant the pachytene stage was not observed. Scale bar represents 5 µm (A,D,G).

### DMC1-binding PhePP domain in exon 14 of Brca2

The meiotic phenotype of *Brca2*^*Δ12-14*^ was clearly distinct from that of *Hsf2bp* knockout. It was more severe, and in the females involved complete loss of DMC1 but not of RAD51 recombinase accumulation. This latter observation strongly suggested that the deleted region contains, in addition to the HSF2BP-binding domain, a domain that specifically affects DMC1 function. A previous publication ^16^ suggested that BRCA2 exon 14 encodes a direct BRCA2-DMC1 interaction domain involving the PhePP motif, however, this was not validated in any follow-up biochemical studies, and a specific mutant mouse model testing it revealed no effect on meiosis ^18^. We therefore proceeded to replicate and extend the previously published observation using recombinant human DMC1 and BRCA2 fragments expressed in *E. coli* (Figure 4, S5).

**Figure 4.**
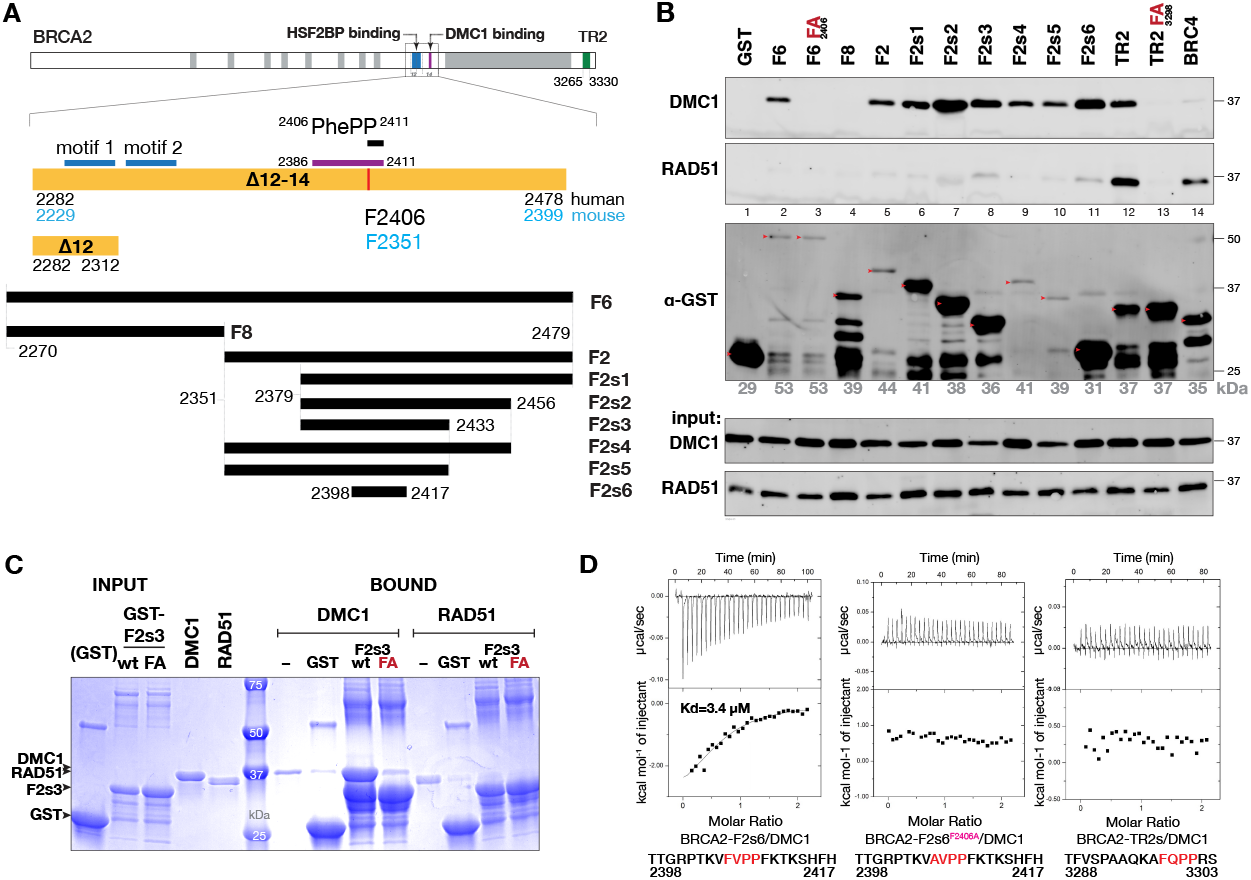
Biochemical characterization of the DMC1-BRCA2 interaction. **(A)** BRCA2 regions deleted in the mice (yellow) and studied in pull-downs (black), HSF2BP-interacting motifs, PhePP motif and the peptide used in ref. 16 are indicated. Residue numbers for human BRCA2 are shown in black, mouse — in blue. **(B)** GST pull-down with the indicated GST-BRCA2 fragments immobilized on GSH-sepharose beads and used to precipitate recombinant DMC1 or RAD51, followed by immunoblotting with anti-RAD51 antibody (cross-reacts with DMC1) and anti-GST antibodies. Full-size GST-fragment bands are indicated by red arrows, predicted Mw listed below the blot. Experiment was performed twice with same results. **(C)** Co-precipitation of purified recombinant untagged DMC1 and RAD51 with purified GST-tagged BRCA2 fragment F2s3 (wild type (wt) and F2406A variant (FA)) immobilized on the beads. Bound proteins were analyzed by SDS-PAGE and stained with Coomassie. **(D)** ITC analysis of the interaction between untagged DMC1 and synthetic peptides corresponding to BRCA2-F2s6 fragment, its F2406A variant, and the peptide from the RAD51-binding TR2 domain, containing a similar FxPP consensus. Experiments were performed twice with the same result, replicates are shown in Figure S5.

DMC1 co-precipitated with the GST-tagged BRCA2 fragment F6, corresponding to the region deleted in our mouse model, and with F2, corresponding to exon 14, but not with F8 corresponding to exons 12-13, and F2406A substitution within the PhePP motif completely abolished this interaction (Figure 4A,B). Using a truncation series, we narrowed down the interaction domain to BRCA2 residues 2379-2433, fragment F2s3. Purified F2s3, but not its F2406A variant, co-precipitated with purified DMC1 but not with RAD51, further demonstrating that the interaction is direct (Figure 4C). We used isothermal titration calorimetry (ITC) to determine that the small synthetic peptide F2s6, corresponding to the conserved region of F2s3, still bound to the purified untagged DMC1 with a Kd of 3.4 µM, and no interaction was observed when the same assay was performed with the F2406A variant. In addition, testing a similar motif encoded by exon 27, in the so-called TR2 region, revealed that despite the presence of a similar (FQPP) motif, that was shown to bind RAD51 38, this fragment does not interact with DMC1 (Figure 4D, S5A). Finally, we confirmed our observations using mouse proteins and the mBRCA2-F2351D substitution (equivalent to human F2406D) introduced in the PhePP-mutant *Brca2* strain ^18^ (Figure S5B). Taken together, these results confirm the presence of a specific DMC1-interacting motif in the region encoded by exon 14 of BRCA2.

### Distinct RPA foci patterns in Brca2Δ12-14 versus wild type zygotene meiocytes

The *Brca2*^*Δ12-14*^ mouse model provides opportunities for studying the initial steps of meiotic DSB repair, due to the complete lack of recombinase loading in spermatocytes. In particular, in this mutant initial RPA loading on single-stranded (ss)DNA should still occur (RPA on resected ends; RPA-R), while possible RPA loading on ssDNA in the D-loop (RPA-D) would be expected to be abrogated in the absence of recombinases. Thus, comparing RPA localization patterns between wild type and *Brca2*^*Δ12-14*^ meiocytes might provide insight in which localization patterns are specific for the initial RPA loading on resected ssDNA. We used the super-resolution microscopy technique SIM to visualize RPA localization in *Brca2*^*Δ12-14*^ and control spermatocytes and oocytes (Figure 5A-B).

**Figure 5.**
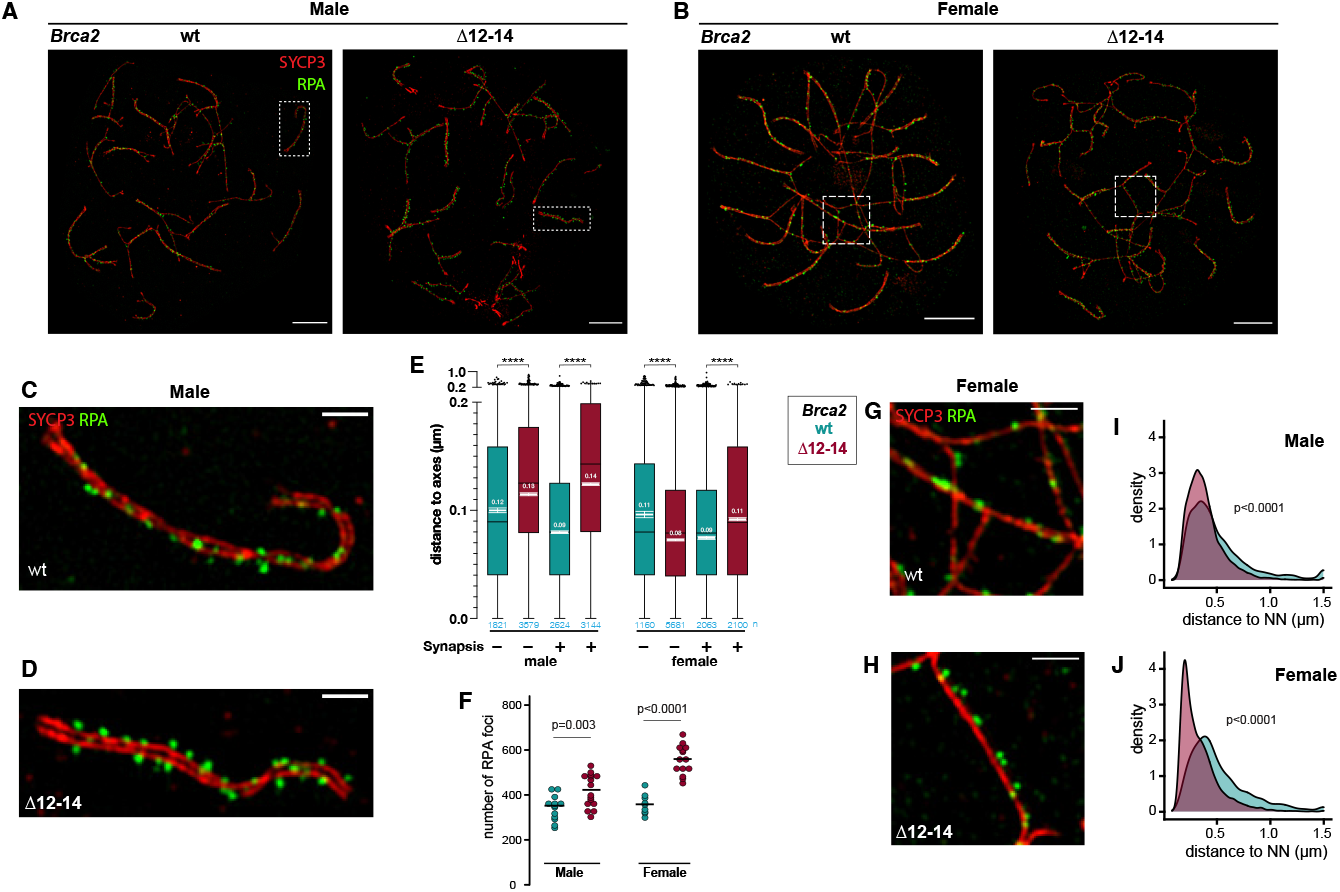
Super-resolution imaging analysis of RPA localization in zygotene meiocytes of *Brca2*^*Δ12-14*^ mice. Structured illumination microscopy imaging of RPA (green) and SYCP3 (red) on nuclear surface spread spermatocytes **(A)** and oocytes **(B)** from wild type and *Brca2*^*Δ12-14*^ mice. **(C, D)** Close-up of SIM image of RPA (green) and SYCP3 (red) on nuclear surface spread spermatocytes from wild type (C) and *Brca2*^*Δ12-14*^ (D) spermatocytes to show the difference in RPA positioning relative to SYCP3. The whole nucleus is indicated in A. **(E)** The shortest distance for each RPA focus to the axial elements (for unsynapsed regions) or to the center of two lateral elements (for synapsed regions) in wild type and *Brca2*^*Δ12-14*^ spermatocytes and E17.5 oocytes at zygotene is plotted as Tukey’s boxplot (median, 25% and 75%) and with mean values (white lines with s.e.m. error bars) overlayed. **(F)** Quantification of the number of RPA foci at zygotene nuclei for male and female *Brca2*^*Δ12-14*^ and control mice. Close-up of SIM image of wild type **(G)** and *Brca2*^*Δ12-14*^ **(H)** E17.5 oocytes to show the difference in paired RPA foci. The whole nucleus is indicated in B. Distribution of nearest neighbor distances for RPA foci of male **(I)** and female **(J)** *Brca2*^*Δ12-14*^ and control mice to show an increase in close-positioned paired RPA foci in *Brca2*^*Δ12-14*^ oocytes. Distance distribution is displayed as kernel density plots produced using R software; distances larger than 1500 nm were plotted as 1500 nm. Scale bar represents 5 µm **(A**,**B)** and 1 µm **(C**,**D**,**G**,**H)**. Statistical significance (p-values) using a Kruskal-Wallis test with Dunn’s multiple comparison **(E)**, two-tailed unpaired t-test **(F)** or Kolmogorov– Smirnov test **(I**,**J)** are indicated in the figures.

First, we investigated the position of RPA foci relative to the axial and lateral elements of the SC by measuring the shortest distance from each focus to the axial elements (for unsynapsed regions) or to the center of two lateral elements (for synapsed regions), in zygotene(-like) nuclei (Figure 5C-E) and found that the mean distance was significantly larger in *Brca2*^*Δ12-14*^ than wild type spermatocytes for both unsynapsed as well as synapsed regions (0.139 µm std.dev.

0.085 vs 0.103 nm std.dev. 0.089, respectively, p<0.0001, synapsed and unsynapsed taken together). It should be noted that for the synapsed regions, we compare homologous (wild type) to nonhomologous (mutant) synapsis. For oocytes there were also clearly significant differences, but the effects were very small; a slightly smaller distance for unsynapsed regions, and slightly larger for synapsis regions, in mutant versus wild type.

Second, we compared the number of RPA foci (Figure 5F). In *Brca2*^*Δ12-14*^ oocytes, we found a median of 558 RPA foci per nucleus, compared to 357 in wild type oocytes (p<0.0001). In *Brca2*^*Δ12-14*^ spermatocytes fewer RPA foci were observed (421), but still more than in wild type spermatocytes (351, p<0.003). This increase in both male and female mutants is consistent with our observations using confocal microscopy (Figure 2E, 3E), but the numbers of detectable RPA foci are higher using SIM than using confocal microscopy. This indicates that some RPA foci were closer than the resolution limit in the confocal images, especially in *Brca2*^*Δ12-14*^ oocytes. Indeed, in the SIM images of mutant oocytes we observed a distinct RPA positioning pattern at unsynapsed regions, with often two RPA foci positioned closely together as a doublet in a more or less side-by-side arrangement along the axial element (Figure 5G,H). These RPA doublets were much less apparent in wild type oocytes or in wild type and *Brca2*^*Δ12-14*^ spermatocytes. To further analyze these doublets, we determined the nearest neighbor distance for all RPA foci, and indeed observed an almost doubled frequency for a distance of ∼200 nm in *Brca2*^*Δ12-14*^ oocytes compared to controls (Figure 5I,J), while the RPA nearest neighbor distance distribution patterns of wild type oocytes and spermatocytes, as well as of *Brca2*^*Δ12-14*^ spermatocytes were all very similar, with a peak at around 350 nm (387 nm for wild type oocytes, 352 nm for wild type spermatocytes, and 318 nm for *Brca2*^*Δ12-14*^ spermatocytes).

### RPA on tensed DNA tethers

The third observation from our SIM analysis were RPA structures, whose distinct morphology and striking absence in the mutant prompted us to systematically characterize them in the wild type. These were highly elongated individual RPA foci or linear constellations of circular foci positioned along an invisible straight line that appears to be connecting two chromosomal axes (Figure 6A, S6). They become detectable in leptotene, where the invisible line they define projects towards distinct kinks in the axial elements, indicating a stretched connection between them. Since RPA bound to ssDNA forms amorphous aggregates and only assumes extended conformation when a DNA-stretching force is applied ^39^, this appearance and the dependence on HR leads to the conclusion that such RPA staining marks a partially single-stranded **tensed DNA tether** between the homologous axes. Consistent with this, antibodies against SPATA22 and BRME1, two other ssDNA-associated proteins ^25,26,28,40^, also recognized similar structures (Figure S7). In early zygotene, tether-associated RPA patterns between completely unsynapsed axial elements with distinct kinks create a characteristic “butterfly” appearance (Figure 6A structures #2-5, and Figure S6C) that resembles the “pinched” configuration of zebrafish chromosomes initiating synapsis ^41^. Tethers get more numerous as homologues progress towards alignment and synapsis in zygotene, and become similar to the RPA structures reported previously by others, and often termed bridges ^42–44^. No tethers were observed in the *Brca2*^*Δ12-14*^ spermatocytes (16 nuclei, 2 animals), which shows that their formation/stabilization requires functional HR. We analyzed all 473 tethers we spotted in the 85 nuclei from four different animals in two experiments, measuring the distance between the axes, presence of kinks and chromosomal context in terms of alignment and/or synapsis (Figure 6B,C). The majority (90%) of tethers in a context without parallel alignment (early structure, n=91) displayed kinks. As alignment became apparent, the frequency of kinks reduced to 39% (mid structure, n=197), and in a context of partial synapsis, this percentage was similar (late structure, 35% kinks, n=185, Figure 6C). The length of the inferred tensed DNA tether decreased with prophase progression (Figure 6D) and was smaller when kinks were absent (no kinks: 0.55±0.29 µm, kinks: 0.86±0.41 µm). We noted that in thirteen examples of the fully traceable butterfly configuration we found, axial elements were arranged symmetrically, with longer parts on the same side of the tether (examples #2-#5 in Figure 6A and 9 structures in Figure S6B (#3) and S6C #1,2,3,4,5,9,13,15), and only in two examples the orientation appeared reversed (Figure S6C #7 and S7D #2). This strong preference for a symmetrical orientation in accordance with the expected homology suggests the presence of additional connections between the axes, which constrain axis orientation but are not visible in the staining. We also noted that only 18% (13/73) of early arrangements had multiple RPA-marked tethers, while the majority of mid and late chromosomal arrangements were connected by more than one visible tether: 75% (58/77) and 57%. (52/92), respectively. Together, these findings point towards the hypothesis that tethers and their chromosomal arrangements are HR-specific chromosome connections that contribute to homologous chromosome pairing and synapsis.

**Figure 6.**
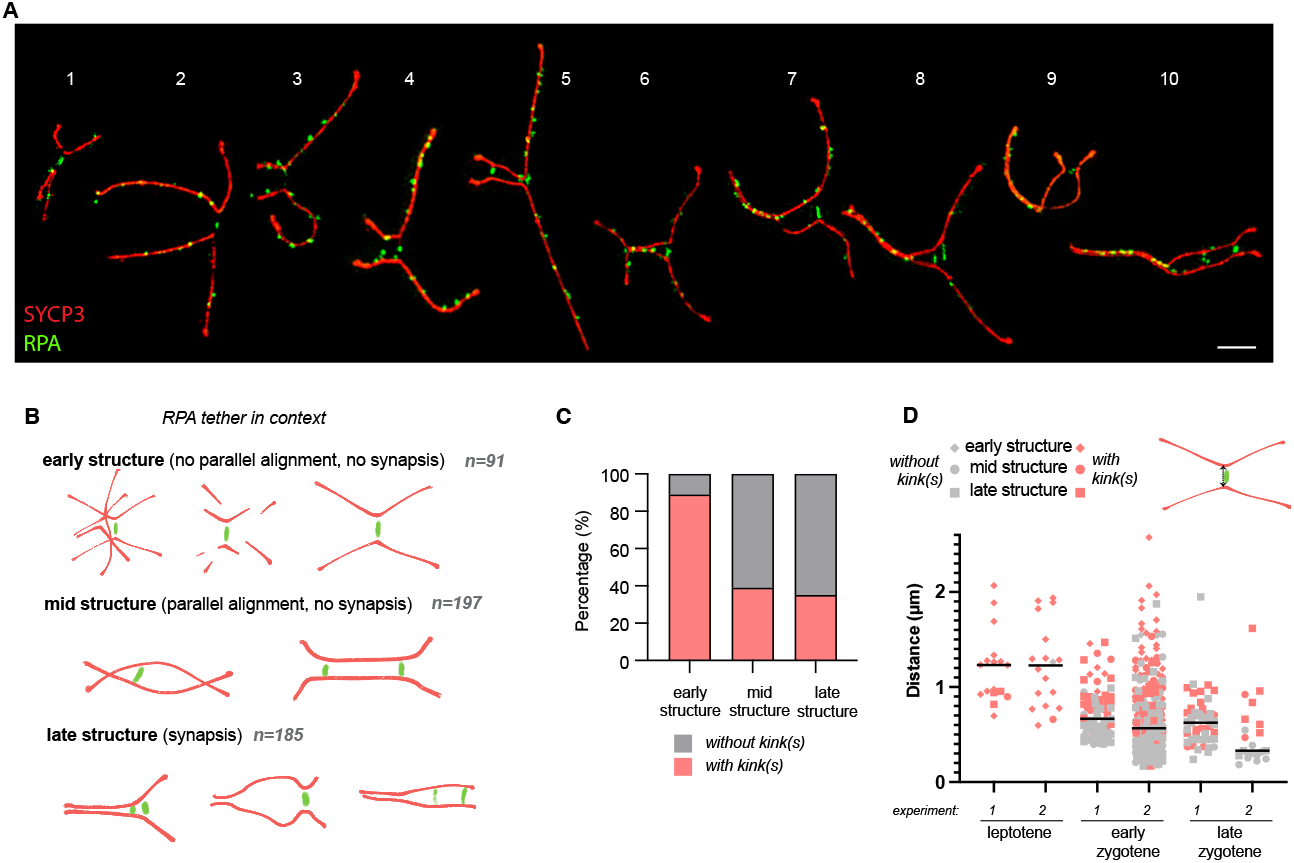
DNA tether-associated RPA foci identified by SIM imaging. **(A)** Compilation of examples of the tether-associated RPA foci observed in SIM data from wild type spermatocytes. RPA is green, SYCP3 is red. These structures are not observed in *Brca2*^*Δ12-14*^ spermatocytes. For the purpose of visualization, images of the chromosome pairs were isolated from the different images of nuclei and pasted together in this combined image. Scale bar 2 µm. **(B)** Several cartoons of possibilities of the different contexts where RPA tethers are present: early (no alignment, no synapsis), mid (alignment, no synapsis) and late structures (synapsis) including the number of RPA tethers in those categories observed in 85 wild type nuclei. **(C)** Quantification of the presence (light red) or absence (grey) of kinks in the chromosomal axes around the RPA tether in the different contexts of the RPA tether. **(D)** Quantification of the distance between the two points on the chromosomal axes to which the line defined by the RPA tether projects per meiotic stage. The presence (light red) or absence (grey) of kinks is indicated. The different structures are indicated by the shape of the points (triangle represents early structure, circle represents mid structure, square represents late structure). Data from two independent experiments is plotted.

## DISCUSSION

### The meiotic domain of Brca2

We generated and characterized a mouse strain that separates the meiotic functions of BRCA2 from its essential role in somatic HR. Deletion of *Brca2* exon 12 to 14 resulted in a complete block of meiotic HR and infertility in both sexes, consistent with reported *Brca2* hypomorphic mouse models ^11,12^ and BRCA2 mutants in other species ^3–10^. As no developmental or other somatic defects were detectable, we propose that the deleted region constitutes the “meiotic domain” of BRCA2 ^45,46^. Characterization of DMC1-binding PhePP domain encoded in exon 14 is particularly significant, as previously published data led to conflicting interpretations ^16,18^. Its functional relevance is now supported by the complete loss of DMC1 foci in both sexes and by the biochemical confirmation of direct interaction between PhePP and DMC1. In a separate study ^47^ we explored this further by analyzing the complex between BRCA2-PhePP and DMC1 by X-ray crystallography, showing that PhePP stabilizes DMC1 filaments, and expanding the observations to other similar “P-motifs”, and consistent data has been published by others ^48,49^.

### Sexual dimorphism in RAD51 loading

Based on the two lost domains, *Brca2*^*Δ12-14*^ could be viewed as a sum of *Hsf2bp* and *Dmc1* knock-out phenotypes: complete loss of DMC1 foci and infertility in both sexes as in *Dmc1*^*-/-*32,33^ and reduced RAD51 foci as in *Hsf2bp*^*-/-*19,21,25^. However, the phenotype is more severe: in *Brca2*^*Δ12-14*^ spermatocytes RAD51 foci are completely undetectable, while still present, at least to some extent, in each of two knock-outs. This suggests a functional interaction between the two deleted domains, presence of other functional domain(s) in the deleted region, or unanticipated effects of the deletion on the adjacent BRCA2 domains. Furthermore, no meiotic defects in the previously described *Brca2-F2351D* strain also challenges the straightforward “sum of knock-outs” interpretation, as this substitution completely abolished DMC1-PhePP interaction in our *in vitro* experiments (Figure S5B). It is possible that *in vitro* experiments exaggerate the effect of the point mutation on the interaction, and that in meiocytes BRCA2-F2351D still interacts with DMC1. We previously observed such discrepancy for HSF2BP-BRCA2: while *in vitro* the interaction of the mutant BRCA2 (*Δ*12) with HSF2BP was reduced >100 fold or completely abolished, in testis the reduction was only up to 2 fold ^22^. Detailed side-by-side re-analysis of the two strains and possibly creation of additional mouse strains with a specific deletion of the PhePP domain are required to address this question further. In a striking contrast to spermatocytes, RAD51 foci are present in *Brca2*^*Δ12-14*^ oocytes and in zygotene even remain more persistent than in the wild-type. This highlights a sexual dimorphism in RAD51 loading, suggesting that in oocytes HSF2BP binding may not be absolutely required to localize BRCA2 to DSBs, or that RAD51 loading may be partially BRCA2-independent. Both HSF2BP and BRME1 still associate with DSB sites, but notably HSF2BP foci numbers were reduced while BRME1 numbers and intensity were increased. This suggests that BRCA2 affects HSF2BP localization, and that BRME1-HSF2BP association is not constitutive.

### DSB formation stimulates synapsis

We observed more complete end-to-end synapsis in *Brca2*^*Δ12-14*^ than in *Spo11* knockout spermatocytes, but it was between non-homologous chromosomes. Since repair is arrested in the mutant, this shows that DSB induction and possibly end resection by themselves somehow stimulate synapsis, albeit nonhomologous. Synaptic adjustment mechanisms, as often observed in cases of non-homology due to translocations, duplications, or inversions ^50,51^, could contribute to the complete-looking synapsis. Since partially synapsed chromosome pairs in *Brca2*^*Δ12-14*^ spermatocytes always have synapsed telomeres (Figure S3F,G), while a third of wild type chromosomes do not, we hypothesize that non-homologous synapsis initiates at telomeres, while homologous synapsis can begin interstitially. Possible explanations to increased synapsis in the presence of DSBs could include modulation of chromosome movements by DSB formation ^52–55^ or existence of DSB-dependent and independent components in the SC assembly mechanism ^56^.

### RPA doublet formation in Brca2Δ12-14 oocytes

Previously, it has been suggested that RPA foci in-between synapsed regions represent post-invasion structures (D-loop; RPA-D foci), while pre-invasion RPA-R foci on resected ssDNA are associated with axial elements or the outer edge of SC ^42–44,57^. As expected, RPA-D foci were lost in our HR mutant. However, the total RPA foci number increased. Most of this is likely due to continued DSB formation in the absence of repair and homologous synapsis, but the distinct 200 nm-apart doublets accounting for the additional increase in foci numbers in oocytes (Figure 5J) calls for other explanations: RPA foci numbers will depend on whether one or both DSB ends are resected and whether the distance between them exceeds microscopy resolution limit. The simplest interpretation to the oocyte-specific doublets is that one (or both) of these parameters is different from spermatocytes. This may be related to the partial retention of RAD51 function in oocytes or reflect a more general mechanistic difference in meiotic HR between sexes ^58^. The peak of ∼ 200 nm in the distance distribution could reflect structural properties of a protein complex, e.g. the end-tethering RAD50 ^59,60^. The alternative, but in our view less likely explanation, is that the doublets represent exactly two DSBs in a close and constrained proximity, either on the same chromatid or on the sister, like the recently described double-DSBs ^61,62^.

### The tensed DNA tether model of homologous synapsis

Our observations indicate the existence of HR-dependent DNA tethers, with partially single-stranded regions marked by RPA, some of which appear tensed on chromosome spreads. Elongated proteinaceous structures connecting co-aligned homologs have been documented previously ^42–44,57,63–66^, including RPA foci linking synapsed or near-synapsed homologs in mice and humans ^42–44,57^, sometimes associated with kinks (“inflections” ^44^) in the axial elements, and bridges enriched in ZMM proteins between *Sordaria* homologs co-aligned at 400-200 nm ^65,66^. However, the earlier and less frequent structures at larger distances (1-12 µm) that we report here have largely eluded attention, unless associated with clearly co-aligned chromosomes as observed in different plants ^63,67^ and more recently, in human ^42^. The reduced tether length with the progression of meiotic prophase I we observed requires a mechanism for reeling in the connecting DNA. Molecular motor activity (e.g. cohesin loop extrusion^68^), a Brownian ratchet^69,70^ and condensation of tether-associated proteins^65,71^ are possible mechanisms. From the symmetric orientation of the “butterfly” configuration (Figure 7A), we infer that there are more tethers than can be detected via tension and RPA staining. Although clustering of centromeres/proximal telomeres of the acrocentric chromosomes in mice ^72–74^ may also contribute to this symmetry, the increasing detection of the HR-dependent tethers in zygotene when HR initiation is over, further supports the untensed orientation-constraining tethers undetected by our stainings. This led us to two hypotheses resulting in the “tensed DNA tether model” of homologous synapsis described below (Figure 7, S8, Supplementary Movie).

**Figure 7.**
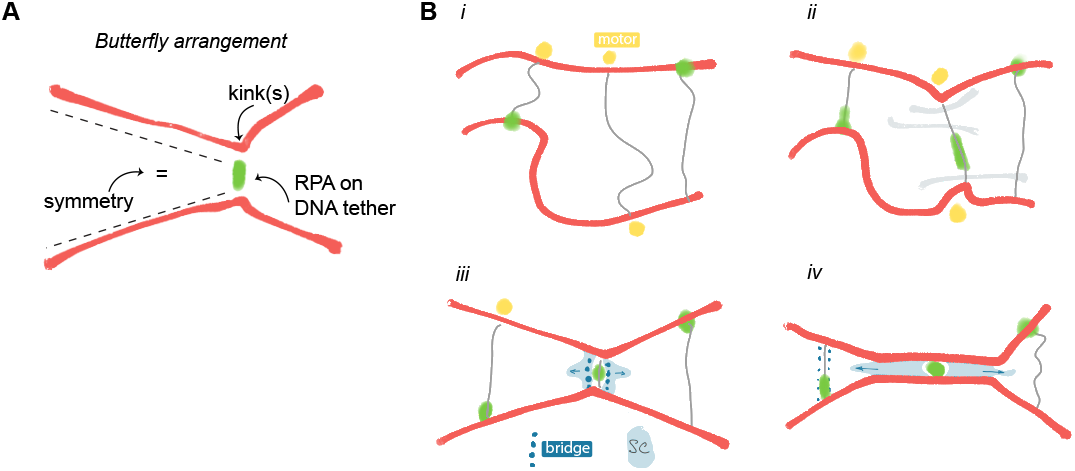
Tensioned DNA tether model. **(A)** Key features found in the butterfly arrangement. **(B)** The tensioned DNA tether model of homologous synapsis, motor scenario (see Figure S8, S9 and the Supplementary Movie for additional scenarios). Grey lines indicate DNA connections between homologs, yellow circles indicate the hypothesized molecular motors that reduce the length of the DNA tethers bringing homologs closer together (step *i*). Occasionally, clashes with other chromosomes (light grey, step *ii*) create counter force sufficient to induce mechanical tension that results in RPA recruitment. Kinks in the axial elements may result from spreading of chromatin on glass, or might also form in the cells and as such propagated upon spreading. As the homologs come closer together at one or multiple points, proteinaceous connections (bridges and eventually synapsis) can form, stabilizing the interaction (step *iii*). Once initiated, synapsis rapidly progresses, while continued shortening of the tethers can facilitate bringing together unsynapsed regions (step *iv*).

The first hypothesis, essential for the model, is that DNA tether shortening does not simply accompany the homolog co-alignment but is the critical mechanism for holding and/or pulling homologs together. Similar ideas have been suggested decades ago to explain presynaptic events in *S. cerevisiae* and *Sordaria* ^75–78^, and the tethers we describe during meiotic prophase progression provide the first quantified morphological evidence supporting this hypothesis for mammals. We further argue that this hypothesis is unavoidable to bridge two gaps between HR and synapsis. The first gap is temporal: recombinase loading and HR are initiated in early leptotene, when fragments of axial elements only begin to form, while co-alignment and synapsis happen in zygotene, and require complete continuous axes. Preserving the precious HR contacts through the turbulent events of leptotene–zygotene transition (axis assembly, telomere clustering, chromosome motions and collisions) requires robust flexible connections, and HR-dependent dynamic DNA tethers are the best if not the only candidate for the role. The second gap is spatial: between the site of homologous DNA contacts and the corresponding “homologous” regions of the axes that will engage in protein-protein contacts mediating synapsis (Figure S8D). Donor DNA is in the chromatin loop and can be micrometers away from its axis attachments site, which does not differ from any other (non-homologous) loop attachment site. The ends-apart model also proposes that the end searching for homology is also micrometers away from its axis attachment site. Clashes between the axial elements per se, without coordination with the off-axis HR contact, are likely to occur but would be nonproductive. However, they may initiate non-homologous synapsis, in particular in cases of HR problems, as observed in our (Figure S3H) and other HR mutants ^11,32,33,79–81^. While further molecular specification of the model would be purely speculative at this point (Figure S9), DNA tether shortening through loop extrusion by axis-associated cohesins seems one plausible mechanism for bridging the spatial gap, providing a molecular term to the “axis capture step 3” proposed by Zickler and Kleckner in 1999 ^64^ , and subsequent suggestions of “reeling in” the homolog at a somewhat later stage ^75–78,82^ (Figure S8C). Intriguingly, one of the consequences of the meiotic cohesin Rec8 knock-out in mouse is distant homolog co-alignment 83, which is the phenotype expected if both tether shortening and non-homologous synapsis were blocked.

The second hypothesis is that tether tensioning (a) can happen in cells and (b) is functional. Regarding supposition (a), chromosome spreading procedure has been our default explanation for the tether-revealing axial kinks and RPA foci elongation. Furthermore, we believe that the spreading protocol used in our laboratory (using ample fixative on the slide when applying the cell suspension) facilitates unambiguous detection of the tethers, as it maximizes separation of chromosomes. This does not mean that all of the tether-defining structures are artefactual. First, the appearance of spread chromosomes does reflect their endogenous structure, as exemplified by the typical and reproducible shape of the XY body in pachytene. Second, vigorous chromosome movement can tension even a static tether. Third, reeling in of the tether does require some tension, which may become extreme when motion is impeded by chromosome entanglement. Fourth, such extreme tensioning in the nucleus can explain the origin of ssDNA and RPA within the tether. It is unlikely that they represent off-axis HR contact or the recombinosome ripped off the axis by spreading, since we do not observe recombinases in these foci, only the ssDNA-associated proteins (RPA, SPATA22, BRME1). In addition, studies using optical tweezers revealed that tensioning dsDNA produces ssDNA regions that can bind RPA in a linear arrangement ^84,85^. Similar events are proposed to explain RPA localization to tensioned DNA in ultrafine anaphase bridges ^86^. Fifth, since the role of RPA is to prevent ssDNA (re)annealing, once produced, ssDNA regions will remain RPA-bound after the tension is released, so elongated RPA foci may be result from spreading but still be a vestige of endogenous tension. As to the supposition (b) about functional relevance, tension could be used to trigger a signal (Figure S8E-G) that a donor loop became engaged in an HR intermediate and should be reeled in: once engaged, the loop cannot move unconstrained, comes under tension due to chromosome movement or extrusion, the tension is sensed at the attachment point, changing the directionality ^87^ or increasing the speed of the molecular motor, resulting in a continuous pull. Intriguingly, observations of homolog dynamics during co-alignment in yeast described while this study was under revision suggested that tension may trigger the rapid juxtaposition step ^88^

Taken together, our observations support and extend the model first proposed two decades ago ^76^: HR results in the formation of multiple DNA tethers between the homologs in leptotene, which are somehow reeled in (e.g. meiotic cohesins), capturing homologous contacts and converting them into co-alignment, guiding the formation of bridges and then homologous synapsis.83 Viewing chromatin loops as dynamic, regulated, adjustable structures, also provides straightforward explanations for other directed diffusion events^57,70^ during meiosis as illustrated in Figure S8E.

## METHODS

### Animals

*Brca2*^*Δ12-14*^ mice were generated by two CRISPR/Cas9 cut excision, as described before ^21,22^. Female donor mice (age 5 weeks, C57BL/6 OlaHsd from Envigo) were superovulated by injecting 5-7.5 IE folligonan (100-150 μl, IP (FSH hormone; time of injection ± 13.30 h; day -3). Followed at day -1 by an injection of 5-7.5 IE chorulon (100-150 μl, IP (hCG hormone; time of injection 12.00 h). Immediately after the chorulon injection, the females were put with fertile males in a one to one ratio. Next day (day 0), females were euthanized by cervical dislocation. Oviducts were isolated, oocytes collected and injected with ribonucleoprotein complexes of S.p.Cas9 3NLS (IDT cat. no. 1074181), crRNA and tracrRNA (both with Alt-R modifications, synthesized by IDT). Target sequences for crRNA were TAATATTCCAACCCTCGTGT (*Brca2* intron 11) and TGAGGCTTCCCTTAGGATTG (*Brca2* intron 14). For ribonucleoprotein formation equal volumes (5 µL) of crRNA and tracrRNA (both 100 µM in IDT annealing buffer) were mixed, heated to 95 °C for 5 min and allowed to cool on the bench and frozen. Two injections were performed. On the day of injection, the annealed RNAs (1.2 µL, 50 µM) were thawed, mixed with Cas9 (10 µl diluted to 200 ng/µl in the DNA microinjection buffer (10 mM Tris-HCl, pH 7.4, 0.25 mM EDTA in water) at the final concentrations 0.12 µM Cas9, 0.6 µM of each of the two crRNA:tracRNA complexes in microinjection buffer. Foster mother (minimum age 8 weeks) were set up with vasectomized males in a two to one ratio. Next day, pseudopregnant female (recognized by a copulation prop) were collected. For transplanting the injected oocytes, pseudopregnant female was anesthetized by an IP injection of a mix of Ketalin (12 mg/ml ketamine in PBS)-Rompun (0.61 xylazine mg/ml PBS) 100 μl per 10 g bodyweight). Post-surgery pain relief was given when the mouse was anaesthetized (S.C. Rimadyl Cattle, 5 mg/ml in PBS, dose 5 μg/g mouse). Transplantation resulted in 10 pups from two litters, of which two (one male one female) contained the deletions in the targeted region, as determined by PCR genotyping followed by cloning and sequencing of the PCR product. Another female contained an inversion of the region flanked by the CRISPR/Cas9 cuts; this was not used to establish a colony. Multiple primer combinations were tested during initial genotyping, but mB2i11-F1 AGCTGCCACATGGATTCTGAG, mB2i14-R1 ACATGCAGAGAACAGGGAGC, and mB2e12-R1 GCTTTTTGAAGGTGTTAAGGATTTT, were used routinely. The experimental cohort was eventually formed through back-crossing and inter-crossing from two founders, with the deletion produced by near-direct ligation between the two Cas9 cuts. Routine PCR genotyping of was performed using MyTaq Red mix (Bioline) and using the mentioned primers in 2:1:1 combination, for simultaneous amplification of the wild type and the *Δ*12 alleles (PCR products 663 and 770 bp, respectively).

Testes and ovaries were isolated from mice aged 6 to 32 weeks. Ovaries were also isolated from E15.5 embryos for collection of early meiotic prophase nuclei (leptotene, early zygotene for wild type and leptotene for mutant), and from E17.5 embryos for collection of late meiotic prophase nuclei (late zygotene and pachytene for wild type and early zygotene and late zygotene for mutant).

### Periodic Acid Schiff (PAS) staining of mouse testes

Sections of paraffin-embedded gonads were mounted on slides. Paraffin was melted at 60°C for 60 min, and removed through three 5-minute washes of the slides in xylene. Subsequently, the slides were washed three times in 100% ethanol for 5 minutes. Next, the slides were incubated in 0.5% periodic acid solution (Sigma-Aldrich 395132) for 5 minutes, and rinsed in distilled water. The slides were then incubated in Schiff reagent (Sigma-Aldrich, 3952016) for 15 minutes to stain the tissue, followed by washing the slides in lukewarm, streaming tap water for 5 min. Subsequently, the slides were counterstained with Hematoxylin solution (Klinipath 4085.9005) for 4 minutes and washed in streaming tap water for 5 minutes. The slides were embedded in Glycergel Mounting Medium (Dakocytomation, C0563) (heated to 45°C) and covered with a coverslip.

### Hematoxylin-Eosin staining of mouse ovaries

Sections of in paraffin-embedded gonads were mounted on slides. Paraffin removal was performed as described above, and after two washes in 95% ethanol and one wash in 70% ethanol, the slides were stained with hematoxylin solution (Klinipath, 4085.9005) for 3 minutes and washed in streaming tap water for 5 minutes. Subsequently, the slides were stained for eosin Y (Sigma-Aldrich, HT110232) for 3 minutes, rinsed in tap water for 30 seconds, and dehydrated through 5 minutes 95% ethanol, two washes for 5 minutes 100% ethanol and two washes for 5 minutes xylene. The slides were embedded in DPX Mounting Medium (Merck, HX74511279) and covered with a coverslip.

### Cell culture and clonogenic survival assay

*Brca2*^*Δ12-14*^ and *Brca2*^*Δ12*^ mES cells were engineered as described before ^22^, but using *Brca2*^*GFP/GFP*^ cell line characterized in our previous publications ^89,90^ as the starting point instead of the *Hsf2bp*^*GFP/GFP*^ cell line. mES cells were grown on gelatinized dishes in (0.1% gelatin in water) as described before ^91^ at atmospheric oxygen concentration in media comprising 1:1 mixture of DMEM (Lonza BioWhittaker Cat. BE12-604F/U1, with Ultraglutamine 1, 4.5 g/l Glucose) and BRL-conditioned DMEM, supplemented with 1000 U/ml leukemia inhibitory factor, 10% FCS, 1x NEAA, 200 U/ml penicillin, 200 µg/ml streptomycin, 89 µM β-mercaptoethanol. Clonogenic survival assays were performed in 6-well plates in technical duplicates. Two independent clones that genotyped homozygous for the deletion allele were used in each experiment and considered biological replicates. The experiment was performed three times (n=3 for the wild-type and n=6 for the mutant genotype). Untreated control wells were seeded with 200 cells per well in 2 mL media. Higher seeding densities of 800 and 2000 cells per well were used in the wells treated with higher drug concentrations. One day after seeding, the drugs were added: mitomycin C (MMC, Sigma, M4287-2MG), cisplatin, olaparib, talazoparib (BMN-673, Axon medchem, #2502). After 2-hour (MMC) or overnight (cisplatin, talazoparib) incubation, drug-containing media was removed, wells were rinsed with PBS and refilled with 2 mL fresh media. Colonies were stained (0.25% Coomassie briliant blue, 40% methanol, 10% acetic acid) on day 7 after seeding. Plates were photographed using a digital camera, images were analyzed using OpenCFU software ^92^ to quantify the colonies.

### Meiotic spread nuclei preparation and immunocytochemistry

Nuclei of spermatocytes and oocytes were collected from gonads and spread as described previously ^93,94^ and stored at -80 °C. For immunocytochemistry, the protocol was described before ^22^. In short, slides were washed with PBS, blocked, and incubated with primary antibody overnight. Subsequently, after washing with PBS, the slide was again blocked and incubated with secondary antibody for 2 hours. Finally, the slides were washed with PBS and embedded in Prolong Gold with DAPI (Invitrogen, p36931). The following primary antibodies were used: mouse anti-SYCP3 (1:200, Abcam ab97672), guinea pig anti-SYCP3 ^95^(1:200), guinea pig anti-SYCP2 ^96^(1:100), mouse anti-DMC1 (1:1000, Abcam ab11054), rabbit anti-RAD51 ^97^(1:1000), rabbit anti-BRME1 ^25^ (1:100, #2), rabbit anti-HSF2BP ^25^ (1:30, #1), rabbit anti-RPA ^98^ (1:50), guinea pig anti-HORMAD2 ^99^ (1:100), rabbit anti-SPATA22 (1:100, Proteintech, 16989-1-AP). The following secondary antibodies were used: goat anti-rabbit Alexa 488 (1:500, Invitrogen A-11008), goat anti-mouse Alexa 555 (1:500, Invitrogen A-21422), goat anti-mouse Alexa 633 (1:500, Invitrogen A-21050), goat anti-guinea pig Alexa 546 (1:500, Invitrogen A-11074).

### Microscopy and meiotic analysis

Immunostained nuclear spreads were imaged using a Zeiss Confocal Laser Scanning Microscope 700 with a 63x objective immersed in oil. Images within one analysis were analyzed with the same gain. Images were analyzed using Image J software^100^ and to analyze the meiotic proteins we used methods as described before ^22^. In short, for all proteins (RAD51, DMC1, BRME1, RPA, SPATA22 and HSF2BP) foci quantification was performed using Image J function “Analyze particles” in combination with a manual threshold. A mask of a synaptonemal complex marker (SYCP3, SYCP2, HORMAD2) was used to reduce detection of background signal. Particles smaller than 0.0196 µm^2^ and larger than 0.98 µm^2^ were excluded, except for HSF2BP where only particles smaller than 0.0294 m^2^ were excluded. For HSF2BP, we also adjusted the particle count because of the variable HSF2BP intensity influencing the particle count, by taking into account the average size of 0.25 µm^2^ of an HSF2BP particle (for details see ref. ^22^).

Foci intensity for SPATA22, BRME1, RPA, and RAD51 (female only) was determined as follows: per focus, the intensity was measured and all foci values from a single nucleus were then averaged. For HSF2BP, the mean intensity within the SYCP3 mask was measured. All intensities were normalized by the mean intensity of SYCP3.

For RAD51 and DMC1 quantifications in spermatocytes, we re-used images from our previous publication ^22^ for two wild types (wt_mouse2 and wt_mouse3), and we also generated new images. For BRME1 quantifications in spermatocytes, we re-used images from the previous publication ^22^ for one wild type (wt_mouse3), and we also generated new images.

Meiotic sub-stages were classified into leptotene, early zygotene, late zygotene and pachytene, corresponding to observation of short chromosomal axes, longer chromosomal axes with some synapsis, long chromosomal axes with much synapsis and complete synapsed chromosomal axes, respectively.

For each meiotic protein staining, images from 2 or 3 different animals of each genotype were pooled as indicated in the figure legends. The number of analyzed nuclei is indicated in the figures.

### Fluorescent in situ hybridization (FISH)

FISH was combined with immunostaining on meiotic spread spermatocytes. First, the immunostaining of SYCP3 (mouse anti-SYCP3, 1:200, Abcam ab97672, combined with Alexa 555) was performed as described above. Thereafter, the slides were washed 10 min in 2× SSC (pH 7.5) at 55°C, followed by 5 min washing in 2x SSC at room temperature. The slides were dehydrated using an ethanol series: 1× 5 min 70% ethanol, 1× 5 min 90% ethanol, 2× 5 min 100% ethanol and finally the slides were dried for 5 min at room temperature. Next, 15 µl of the probe painting mouse chromosome 13 (X Cyting Mouse Chromosome Painting Probe for chromosome 13 in green, D-1413-050-FI) was applied on the slide and covered with a coverslip and sealed with photo glue. The sample and probe were denatured for 2.5 min at 78°C on a heating plate. The plate was turned off and the slides were left on the plate for 30 min, followed by incubation at 37°C in a humid box overnight. The next day the protocol of the probe supplier for washing and mounting was followed. In short, the photo glue and coverslip were removed, subsequently the slides were incubated with 0.4x SSC (pH 7) at 72 °C for 2 min and then washed with 2x SSC with 0.05% Tween-20 (pH 7) for 30 seconds. Subsequently, the slides were rinsed with distilled water and air dried. Finally, the slides were mounted with Prolong Gold Antifade Mountant with DAPI.

In total, 152 images of *Brca2*^*Δ*^ ^*12-14*^ spermatocytes (from two different animals) were acquired using a confocal microscope (as described above) and manually scored for number of FISH clouds and degree of synapsis. Nuclei for which the number of FISH clouds was unclear, were excluded from further analysis (30 images in total).

The number of fully synapsed chromosomes was quantified manually. STED images of nuclear spreads of *Spo11*^*-/-*^ spermatocytes immunostained for SYCP3 originated from the data we collected for our recent publication ^101^. For *Brca2*^*Δ12-14*^, the SIM images of nuclear spreads of *Brca2*^*Δ12-14*^ spermatocytes were analyzed (see below). In total, 24 *Spo11*^−*/*−^ nuclei and 16 *Brca2*^*Δ12-14*^ nuclei were used for this analysis, both collected from two different animals.

### Expression and purification of DMC1

Construct pAZ379 for bacterial expression of human his-TEV-DMC1 was engineered by subcloning DMC1 coding sequence into pETM-11 vector using Gibson assembly. Rosetta2 DE3 pLysS *E. coli* strain was transformed with the construct and plated on selective (50 µg/mL kanamycin, 30 µg/mL chloramphenicol) LB-agar plates. Ten colonies were used to inoculate 200 mL selective LB media, the culture was grown overnight at 37 °C with shaking, and used to inoculate 9 L selective LB media. The 9 L culture was grown at 37 °C till OD^600^ reached 0.6-0.8, expression was induced by adding IPTG to 0.2 mM final concentration. Cells were cultured for another 3 h at 37 °C, collected by centrifugation and frozen. The pellet was thawed in two volumes of lysis buffer (3M NaCl, 100mM Tris pH7,5, 10% Glycerol, 0.5mM EDTA, 5 mM β-mercaptoethanol, protease inhibitors (Roche)) and sonicated on ice (10 pulses of 10 seconds). The lysate was incubated overnight at 4 °C on a rotator, and then cleared by centrifugation at 22000× rcf for 45 minutes at 4 °C. Cleared supernatant was supplemented with 5 mM imidazole and incubated with washed (500mM NaCl, 25mM Tris pH8, 10% Glycerol, 0.5mM EDTA, 1mM DTT ) Ni-NTA beads for 1h at 4 °C on a rotator. Beads were washed with 20 mM and 40 mM imidazole (two times 10 mL for each wash). Bound protein was eluted with 400 mM imidazole, fractions analyzed by electrophoresis and pooled, supplemented with TEV protease and dialyzed overnight against 2 L of 150mM NaCl, 10% Glycerol, 50mM Tris pH 8, 5mM β-mercaptoethanol. Protein was further purified using 5 mL HiTrap heparine with a 15CV gradient elution from 150 to 1000 mM NaCl in 25mM Tris pH 8, 10% Glycerol, 1mM DTT. Fractions were pooled and diluted to 100mM NaCl and loaded on a 1ml CaptoQ column. Protein was eluted with a 10CV gradient from 100-1000mM NaCl in 25mM Tris pH 8, 10% Glycerol, 1mM DTT. Fractions were pooled and aliquots were frozen in liquid nitrogen and stored at -80°C.

### Expression and purification of RAD51

The human RAD51 was purified from *E. coli* as previously described ^102–104^. Briefly: The cell pellet containing expressed untagged RAD15 was thawed in two volumes of lysis buffer (3M NaCl, 100mM Tris pH7,5, 10% Glycerol, 4mM EDTA, 20 mM β-mercaptoethanol, protease inhibitors (Roche)) and sonicated on ice (10 pulses of 10 seconds). The lysate was incubated overnight at 4 °C. To cleared supernatant ½ volume of 4M ammonium sulphate was slowly added while stirring on ice for 1hr. After spinning down, the pellet was dissolved in buffer and dialysed overnight. Spin down again and dissolve pellet in buffer (600mM NaCl, 25mM Tris pH7,5, 5% Glycerol, 1mM DTT) for 2hr. Dilute slowly to 200mM and load on 5ml Hitrap Heparine column. Elute with gradient to 1M NaCl. Further purify with Superdex 200 16/60 and final 1ml Hitrap Heparine column.

### Expression and purification of GST-BRCA2 F2s3

Constructs pAZ388 (wt) and pAZ404 (F2406A) for bacterial expression of human his-GST-TEV-BRCA2 F2S3 (aa2379-2433) were engineered by subcloning BRCA2 coding sequence into pETM-30 vector using Gibson assembly. Rosetta2 DE3 pLysS E. coli strain was transformed with the construct and plated on selective (50 µg/mL kanamycin, 30 µg/mL chloramphenicol) LB-agar plates. An overnight culture of a few colonies was grown at 37 °C. 3 liter culture was inoculated with overnight culture and grown at 37 °C till OD600 reached 0.5-0.7, expression was induced by adding IPTG to 0.2 mM final concentration. Cells were cultured for another 3 h at 37 °C, collected by centrifugation and frozen. The pellet was thawed in an equal volume of lysis buffer (1M NaCl, 25mM Tris pH7,5, 5% Glycerol,1mM DTT, protease inhibitors (Roche)) and sonicated on ice (10 pulses of 10 seconds). The lysate was cleared by centrifugation at 22000rcf for 45 minutes at 4 °C. Cleared supernatant was supplemented with 10 mM imidazole and loaded on a 5ml Histrap HP (Cytiva) column, prewashed with buffer buffer A (500 mM NaCl, 25 mM Tris pH7,5, 5% Glycerol, 10mM Imidazol, 1 mM DTT). Wash 3CV with buffer A and elute with 5CV buffer B (500 mM NaCl, 25 mM Tris pH7.5, 5% Glycerol, 250mM Imidazol, 1mM DTT). Fractions were collected and diluted to final 100mM NaCl, 25mM Tris pH8,8, 5% Glycerol, 1mM DTT and load on 5ml Hitrap Q (Cytiva). Wash 3CV and elute with 5CV gradient to 1M NaCl, 25mM Tris pH8,8, 5% Glycerol, 1mM DTT. Peak fractions were collected and loaded on Superdex 75 16/60 column. Fractions were pooled and aliquots were frozen in liquid nitrogen and stored at -80°C.

### GST pull-down assay with immunoblot detection

Bacterial expression constructs for expression of his-GST-TEV-tagged BRCA2 fragments and their amino acid substitution variants were engineered using Gibson assembly in pETM-30 vector and sequence-verified. Constructs were transformed into Rosetta2 DE3 pLysS *E. coli* expression strain. Two mL of selective LB (50 µg/mL kanamycin, 30 µg/mL chloramphenicol) media was inoculated, the culture was grown overnight at 37 °C with shaking, diluted to 10 mL, further incubated till OD_600_ reached 0.6-0.8, induced with 0.2 mM IPTG, grown for additional 3 h at 37 °C. Cells were collected by centrifugation, pellet was resuspended in 1 mL NETT+DP buffer (NaCl 100 mM, Tris-HCl pH 7.5 50 mM, EDTA pH 8 5 mM, Triton X100 0.5%, freshly supplemented with protease inhibitors (Roche), 1 mM DTT, 1 mM PMSF) and sonicated (10× 5 sec on, 10 sec off). Lysate was transferred to Eppendorf minicentrifuge tubes and cleared by centrifugation in (30 min at 4 °C). Supernatant was mixed with 20 µL GSH-Sepharose beads (GE Healthcare 17-5132-01) and incubated overnight at 4 °C. Beads were collected by centrifugation (500 rcf, 2 min, 4 °C), washed with NETT+ buffer ((NaCl 100 mM, Tris-HCl pH 7.5 50 mM, EDTA pH 8 5 mM, Triton X100 0.5%, freshly supplemented with protease inhibitors (Roche)), and incubated with DMC1 protein solution (∼2 µg in 1 mL NETT+) for 1.5 h at 4 °C. Prior to incubation, bead suspension was vortexed, a 40 µL aliquot was collected as input, mixed with 12 µL 5× sample buffer (50% glycerol, 250 mM Tris HCl pH 6.8, 10% SDS, 0.5% bromophenol blue, 0.5 M β-mercaptoethanol) and denature for 5 min at 95 °C. After incubation with DMC1, beads were washed three times with NETT+ buffer and incubated with 25 µL of Laemmli sample buffer for 5 min at 95 °C to elute bound proteins. Samples (5 µL of eluate) were run on a 13% tris-glycine acrylamide gel, transferred to PVDF membrane, blocked (5% skim milk in PBS-T (PBS with 0.05% Tween-20)) and immunodetected with a mixture of rabbit anti-RAD51 pAb (1:20000 ^97^) and mouse anti-GST mAb (1:5000, B-14 Santa Cruz, sc-138) antibodies overnight. After washes, the membranes were incubated with fluorescently-labelled secondary antibodies (anti-mouse CF680 (Sigma #SAB460199), anti-rabbit CF770 (Sigma #SAB460215)), washed (5 × 5 min PBS-T) and scanned using Odyssey CLx imaging system (LI-COR).

### GST pull-down with protein staining

10µl magnetic glutathione beads (Pierce #88821) were prewashed with buffer (200mM NaCl, 25mM Tris pH7,5, 5% Glycerol, 1mM DTT). 120µg (saturating) of GST or GST-BRCA2 F2S3 was incubated with the beads for 1hr at 4 °C. Beads were washed 3x with buffer. 70µg of DMC1 or RAD51 was added and incubated for 1hr at 4 °C. Beads were washed 3x with buffer. To the beads 40µl buffer and 10µl 5x sample loading buffer was added. Samples were boiled 5min at 90 °C and 10µl was loaded on a 4-20% gradient SDS-PAGE gel (Biorad #4561096). Gel was stained with Coomassie brilliant blue R.

### Isothermal titration calorimetry

Interactions between the DMC1 protein and the synthetic BRCA2 peptides were characterized using a VP-ITC Calorimeter (GE Healthcare). The peptides tested were wild type and F2406A variant of the PhePP (F2s6) fragment and the peptide from the C-terminal RAD51-binding TR2 region (3288-3303) The experiments were performed at 20 °C and duplicated; interactions with F2406A and TR2 were also tested at 10 °C. The buffer was 25 mM Tris buffer, pH 7.5, 100 mM NaCl and 5 mM β-mercaptoethanol. 10 μM of DMC1 in the cell was titrated with 100 μM of BRCA2 peptide in the injection syringe; 10 µL of the syringe volume were injected every 210 s. Data recording were performed using the Origin 7.0 software provided by the manufacturer.

### Structured illumination microscopy (SIM) imaging and analysis

SIM images were acquired on a Zeiss Elyra PS1 system using a 63x 1.40 NA oil immersion objective equipped with an 512x512 Andor Ixon Du 897 EMCCD camera, using five phases and five rotations of the illumination. Diode lasers with a wavelength of 488 and 561 nm were used in combination with an emission filter (BP 495-575 + LP 750, BP 570-650 + LP 750). For a SIM image, a z-stack of 21 images was taken with an optimal interval. Images were taken with a laser power between 2 and 5, exposure time of 100 ms and an EMCCD gain based on strong signal intensity, but without saturation. Reconstruction of the SIM data was performed using ZEN2012 software (Carl Zeiss). To generate a single plane image, the most in focus image was selected from the z-stack.

Coordinate determination of RPA foci and synaptonemal complex in SIM images was performed as follows: the coordinates of the synaptonemal complex were determined by selecting a mask of the axes with a Gaussian Blur (sigma of 2), followed by an automatic threshold (Default dark) which was manually checked and using the Image J function “Analyze Particles” only particles larger than 0.2 µm^2^ were included. The mask was dilated by 238.2 nm and of this mask a selection was created to remove all RPA signal outside of the mask. Using the Image J functions “Skeletonize” and “Points from Mask” the coordinates along the center of the SYCP3 mask were determined. The RPA foci were detected using the Image J function “Find Maxima” with a manually set prominence. Using both coordinates, the shortest distance of RPA foci to the skeletonized center of SYCP3 was calculated. In the violin plot distances larger than 500 nm were plotted as 500 nm. For the nearest neighbor analysis all RPA foci within a nucleus were used and for each RPA focus the nearest RPA focus was determined. The distance between this nearest neighbor pair was plotted in a density plot. Distances larger than 1500 nm were plotted as 1500 nm. For the distance and nearest neighbor analysis, 13 nuclei of wild type zygotene spermatocytes and 16 nuclei of *Brca2*^*Δ12-14*^ zygotene spermatocytes (originating from 2 different animals per genotype) were used and 9 nuclei of wild type zygotene oocytes and 14 nuclei of *Brca2*^*Δ12-14*^ zygotene oocytes (originating from 2 different animals per genotype) were used.

### DNA tether identification and scoring

DNA tethers were inferred from the manual analysis of RPA foci shapes and their localization relative to the SYCP3-stained axial elements. SIM images were inspected independently by two co-authors and the candidate tether-containing regions of interest were cropped out and saved as encoded image files. The presence of a DNA tether was deduced if a straight line could be drawn through an elongated RPA focus or a linear constellation of RPA foci towards the (kinked) chromosomal axes. The context of the tether was scored manually, and was divided into three groups: tethers in a context of an early structure (no alignment and no synapsis), tether in the context of a mid-structure (alignment (parallel orientation of chromosomal axes at the distance larger than synapsis) and no synapsis) and tether in the context of a late structure (synapsis). Of each tether we also indicated the presence of kinks in the chromosomal axes. The presence of kinks was determined based on the degree of deflection of the chromosomal axes around the tether. We analyzed 85 wild type and 16 *Brca2*^*Δ12-14*^ spermatocytes, both from two different animals. The butterfly configuration was characterized based on a tether, kinks in both, complete, chromosomal axes.

### Statistical analysis

For the meiotic data the statistical analysis was performed using GraphPad Prism software (version 9) and R (version 4.0.3). For the confocal foci count the statistical significance was determined using unpaired student t-test comparing wild type with *Brca2*^*Δ12-14*^ in different meiotic stages. For the SIM data, statistical significance was determined using a Wilcoxon test, unpaired student t-test or Kolmogorov–Smirnov test, as indicated in the text or figure legend. The number of analyzed nuclei (collected from 2 or 3 different animals per genotype) is in indicated in the graph, text and/or figure legend.

## Supporting information

Numercial Data plotted in the figures

## ACKNOWLEDGEMENTS

We would like to thank Dr Emmanuelle Martini and Dr Alberto Pendas for the helpful discussions of the model. This study was supported by the Oncode Institute, which is partly financed by the Dutch Cancer Society (KWF). We thank the Josephine Nefkens Cancer Program for infrastructure support. S.M. acknowledges funding by the Institut National de la Santé et de la Recherche Médicale (INSERM). S.M. and S.Z.J. were also supported by funding from the French Infrastructure for Integrated Structural Biology (ANR-10-INSB-05-01).

## DATA AVAILABILITY

Raw microscopy images analyzed in the study have been deposited in BioStudies under accession S-BSST1203. Numerical data plotted in the figures is provided as a supplementary Excel file.

## AUTHOR CONTRIBITIONS

Conceptualization: L.K., S.Z-J., R.K., W.B., and A.Z.; Methodology; L.K., A.M., and A.Z.; Software: L.K.; Investigation: L.K., S.vR-F., S.M. E.S-L., A.M., Y.vL., and J.E., Resources: A.M., and J.E.; Data analyses and interpretation; L.K., S.M., S. Z-J., M.B., R.K., W.B., and A.Z.; Visualization: L.K., W.B., and A.Z; Data curation: L.K., and A.Z.; Writing-Original Draft: L.K., W.B., and A.Z.; Writing-Review & Editing: all authors; Supervision: R.K., W.B., and A.Z.; Funding Acquisition: S.Z-J, R.K

## SUPPLEMENTARY FIGURES

**Supplementary Figure S1.**
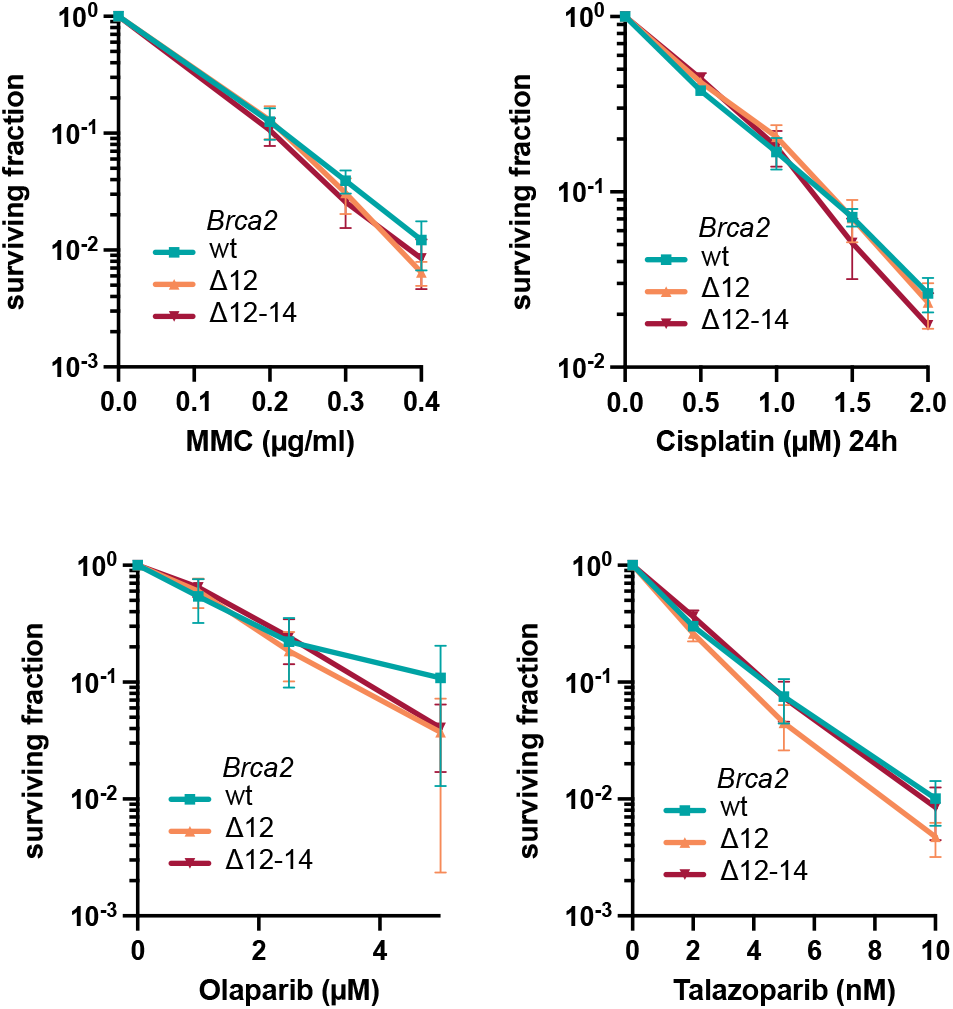
Clonogenic survivals of wild type, *Brca2*^*Δ12*^ and *Brca2*^*Δ12-14*^ mES cells.

**Supplementary Figure S2.**
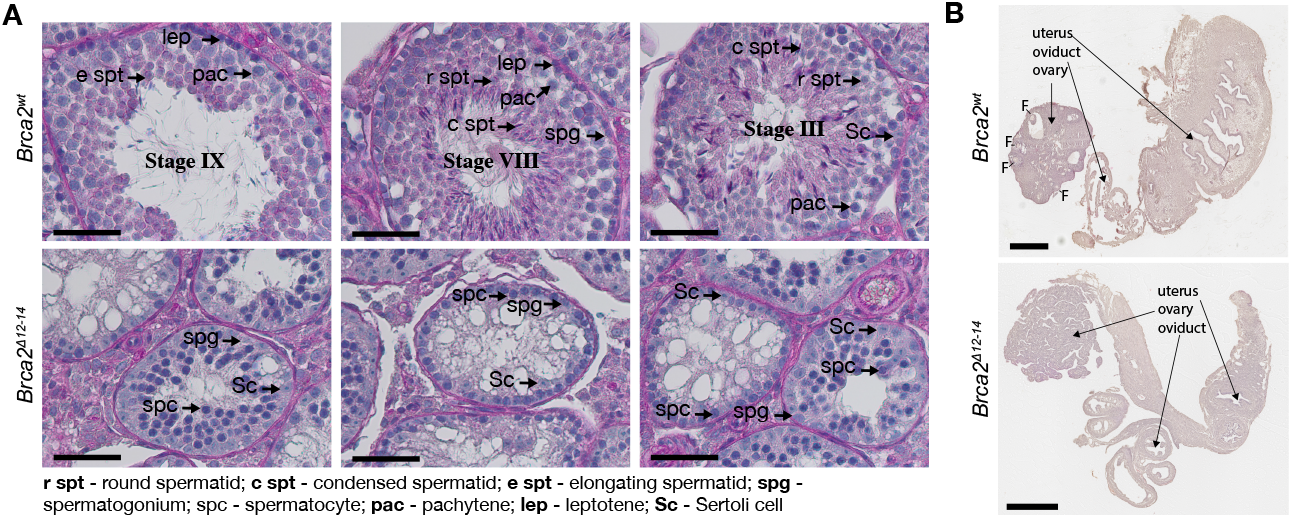
Testis and ovary histology. **(A)** Representative histological images of PAS-staining on testis cross-sections of wild type and *Brca2*^*Δ12-14*^ mice. Seminiferous tubules containing spermatids or spermatozoa (upper panels, wild type) were not present in the mutant, where the development of germ cells proceeded only to the spermatocyte stage (bottom row). **(B)** Representative histological images of HE-staining on ovary cross-sections of 18-week old wild type and *Brca2*^*Δ12-14*^ mice, showing the absence of follicles (labelled F in the wild type) in the mutant. Scale bar represents 50 µm **(A)** and 500 µm **(B)**.

**Supplementary Figure S3.**
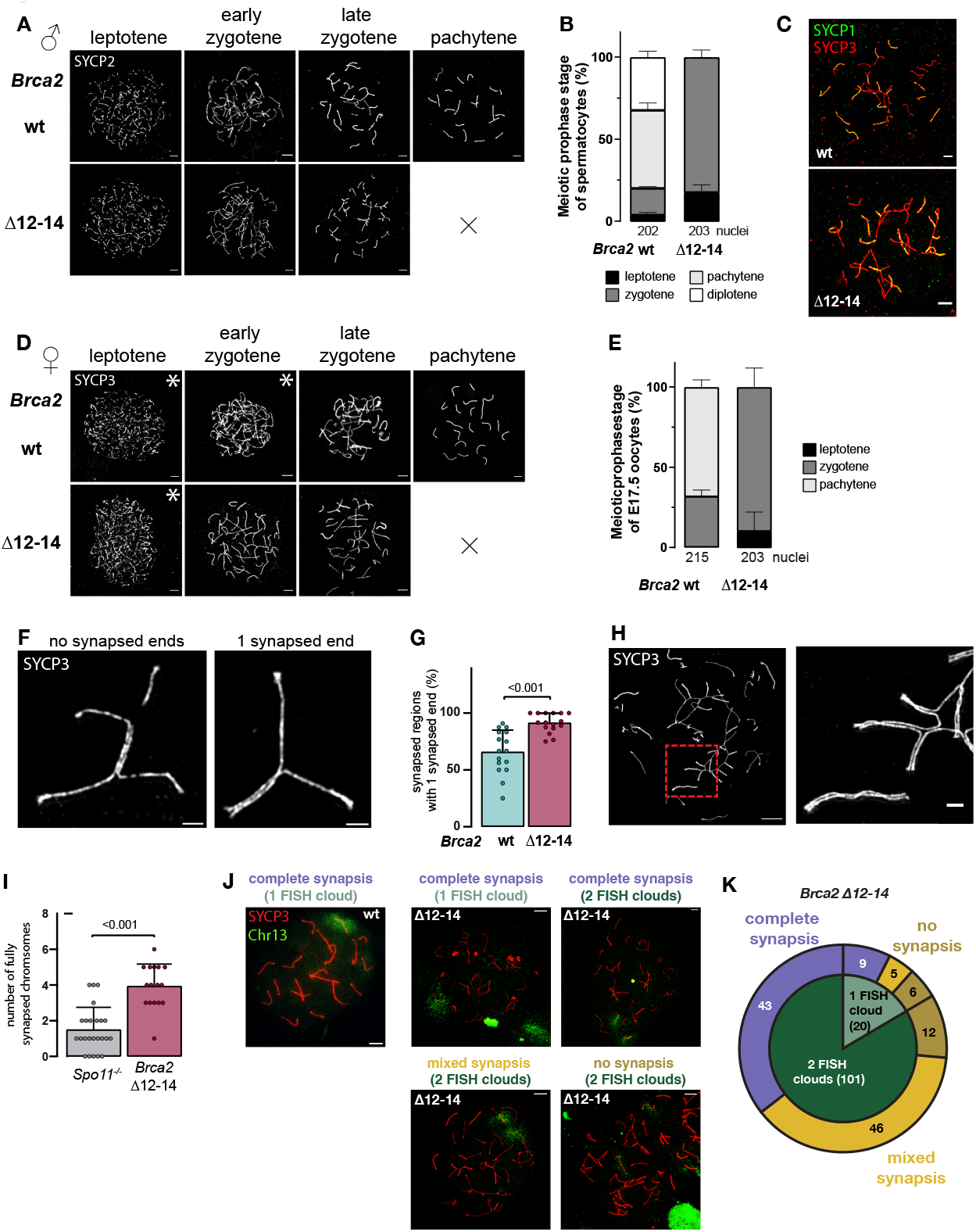
Extended meiotic analysis of synapsis in *Brca2*^*Δ12-14*^ meiocytes. **(A)** Immunostaining for SYCP2 on different meiotic stages of spermatocytes of *Brca2*^*Δ12-14*^ and control mice. **(B)** Quantification of meiotic stages of spread nuclei of spermatocytes of *Brca2*^*Δ12-14*^ and control mice. **(C)** Representative images of spread spermatocyte (wild type and *Brca2*^*Δ12-14*^) nuclei immunostained for SYCP1 (green) and SYCP3 (red). **(D)** Immunostaining for SYCP3 on different meiotic stages of E15.5 and E17.5 oocytes of *Brca2*^*Δ12-14*^ and control mice. Images from E15.5 oocytes are indicated with an asterisk. **(E)** Quantification of meiotic stages of spread nuclei of E17.5 oocytes of *Brca2*^*Δ12-14*^ and control mice. Mutant meiocytes lack the pachytene-stage. **(F)** SIM image of synapsed chromosomal axes in a zygotene wild type spermatocyte immunostained for SYCP3 showing chromosomes with no synapsed ends (left) and one synapsed end (right). **(G)** Bar plot of frequency of synapsed regions in wild type and *Brca2*^*Δ12-14*^ spermatocytes with one synapsed end (fully synapsed chromosomes were excluded). **(H)** SIM image of a whole (left) and zoomed-in (right) late zygotene *Brca2*^*Δ12-14*^ nuclear surface spread spermatocyte immunostained for SYCP3 showing complete and non-homologous synapsis. **(I)** Bar plot of number of fully synapsed chromosomes determined on super-resolution microscopy images of *Spo11*^−*/* −^ and *Brca2*^*Δ12-14*^ zygotene spermatocytes. **(J)** Examples of spread zygotene spermatocytes of *Brca2*^*Δ12-14*^ and control mice immunostained for SYCP3 (red) and combined with fluorescent in situ hybridization of probe for chromosome 13 (green) whereby fully synapsed chromosomes, or chromosomes with mixed synapsis or no synapsis chromosomes were detected and scored for the presence of one or two FISH signals on these structures. **(K)** Quantification of different situations indicated in panel J on 121 spread zygotene *Brca2*^*Δ12-14*^ spermatocytes of two different animals in a pie chart. Scale bars represent 5 µm **(A**,**D, H large, J)** or 1 µm **(F, H small)**.

**Supplementary Figure S4.**
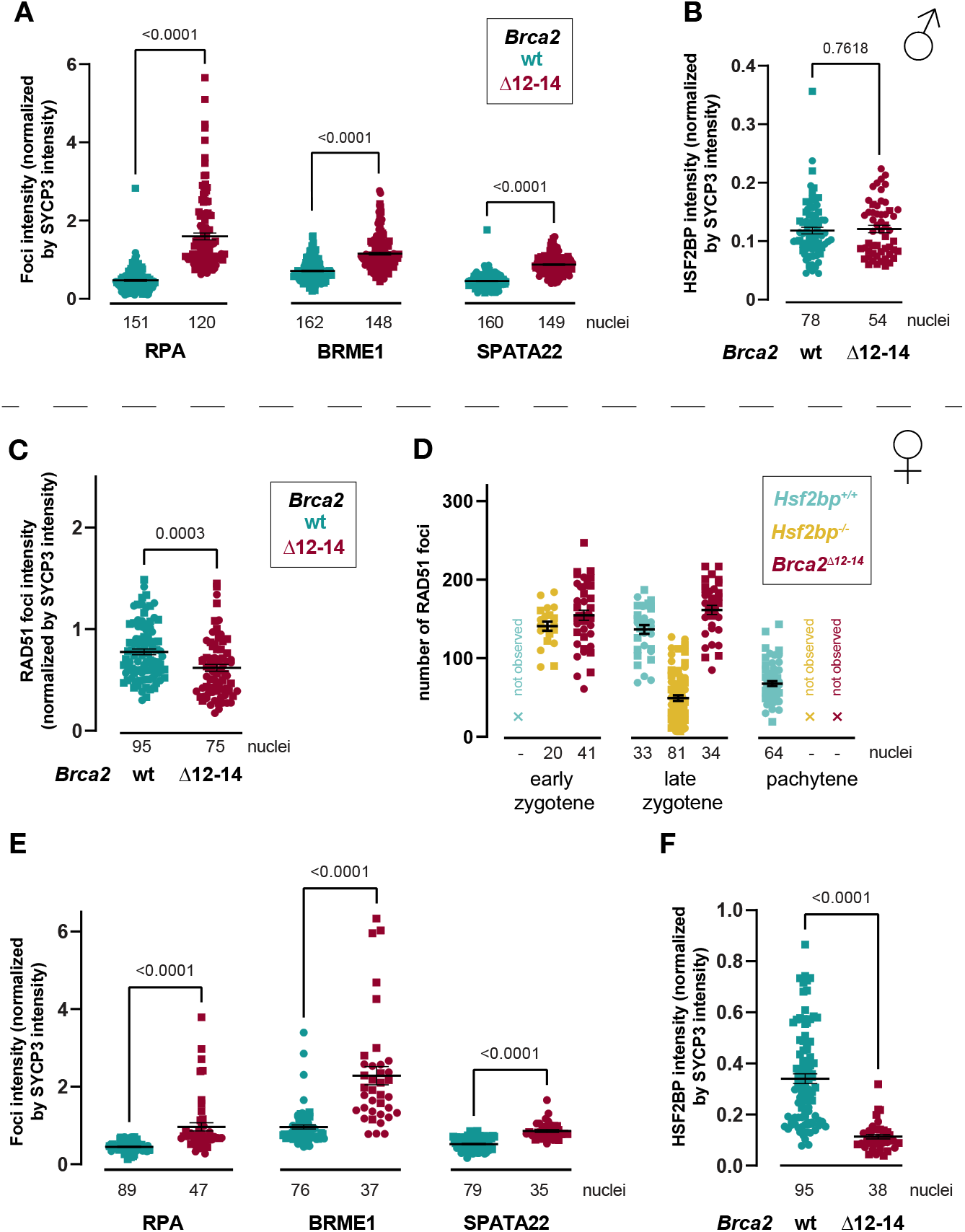
Extended meiotic foci analysis of *Brca2*^*Δ12-14*^ meiocytes. Quantification of RPA, BRME1 and SPATA22 foci intensity in spermatocytes **(A)** and E17.5 oocytes **(E)** of *Brca2*^*Δ12-14*^ and control mice. **(B)** Quantification of HSF2BP intensity in spermatocytes **(B)** and E17.5 oocytes **(F)** of *Brca2*^*Δ12-14*^ and control mice. All intensity quantifications (C-D) were normalized by SYCP3 intensity. Mean, s.e.m., p-values from two-tailed unpaired t-test, and number of analyzed nuclei are indicated in the graphs. Symbol shapes represent individual animals. Number of analyzed nuclei is indicated in the graph pooled from 2 animals per genotype. **(C)** Quantification of RAD51 foci intensity in E17.5 oocytes of *Brca2*^*Δ12-14*^ and control mice. **(D)** Quantification of RAD51 foci number on *Hsf2bp*^+*/*+^, *Hsf2bp*^−*/* −^ and, *Brca2*^*Δ12-14*^ E17.5 oocyte spreads (*Brca2*^*Δ12-14*^ quantification is the same as in Figure 3C). All intensity quantifications **(A-C, E-F)** were normalized by SYCP3 intensity. Mean, s.e.m., p-values from two-tailed unpaired t-test, and number of analyzed nuclei are indicated in the graphs. Symbol shapes represent individual animals. Number of analyzed nuclei is indicated in the graph pooled from 2 animals per genotype. Sea blue color represents *Brca2*^+*/*+^ and burgundy color represents *Brca2*^*Δ12-14*^. Light sea blue and ocher yellow color represent *Hsf2bp*^+*/*+^ and *Hsf2bp*^−*/* −^ respectively. For the mutants the pachytene stage was not observed, and the early zygotene stage was not observed in E17.5 *Hsf2bp*^+*/*+^ oocytes.

**Supplementary Figure S5.**
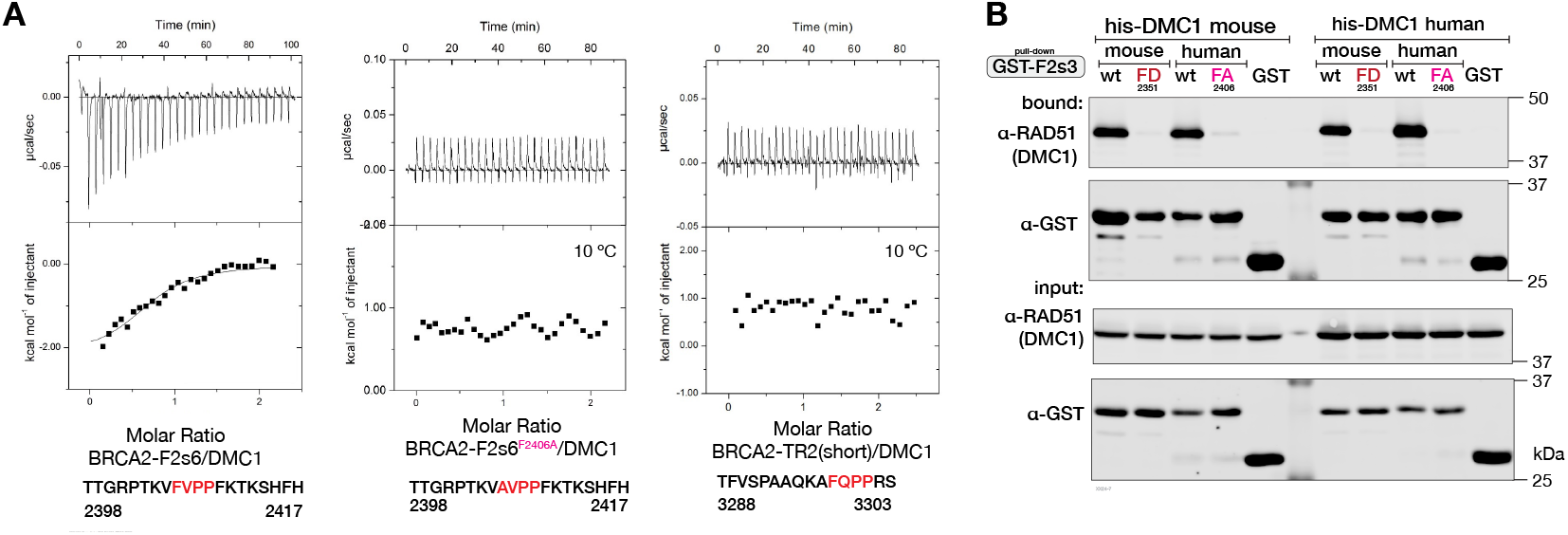
**(A)** Repeat of the ITC experiment shown in Figure 4D. **(B)** GST pull-down using BRCA2-F2s3, the corresponding mouse BRCA2 fragment (I2327-Q2379), their variants with substitutions in the key phenylalanine (F2406A in human BRCA2 and F2351D in mouse BRCA2, to model the mutation introduced in the previously published mouse strain) and mouse or human his-DMC1. Proteins were expressed in *E. coli*, precipitated sequentially form crude lysates and detected by immunoblotting with anti-RAD51 and anti-GST antibodies. The experiment was done twice with the same result.

**Supplementary Figure S6.**
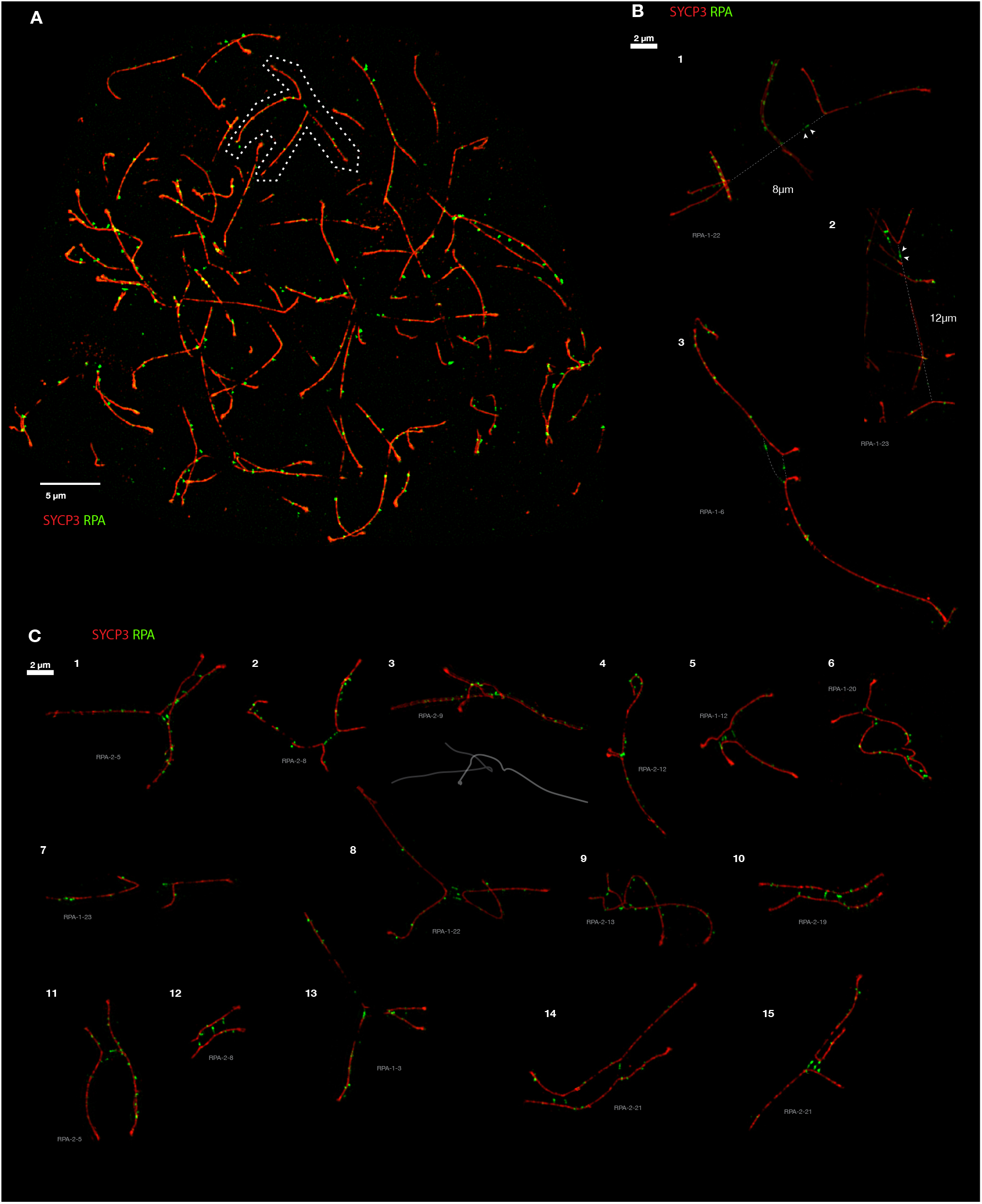
(related to Figure 5). Extended galleries of RPA tethers in the spermatocyte SIM data. **(A)** Illustration of the cropping mask applied to isolate chromosomes linked by RPA tethers for analysis and to produce galleries shown in Figure 5K and panels B and C. The dotted white outline indicates the structure Figure 5K, #2. **(B)** Examples of three structures with exceptionally long RPA tethers that could still be traced. White dotted lines indicate the inferred connections, length is indicated. Image of origin is indicated. **(C)** Extended gallery of RPA tethers. Scale bar = 2 µm.

**Supplementary Figure S7.**
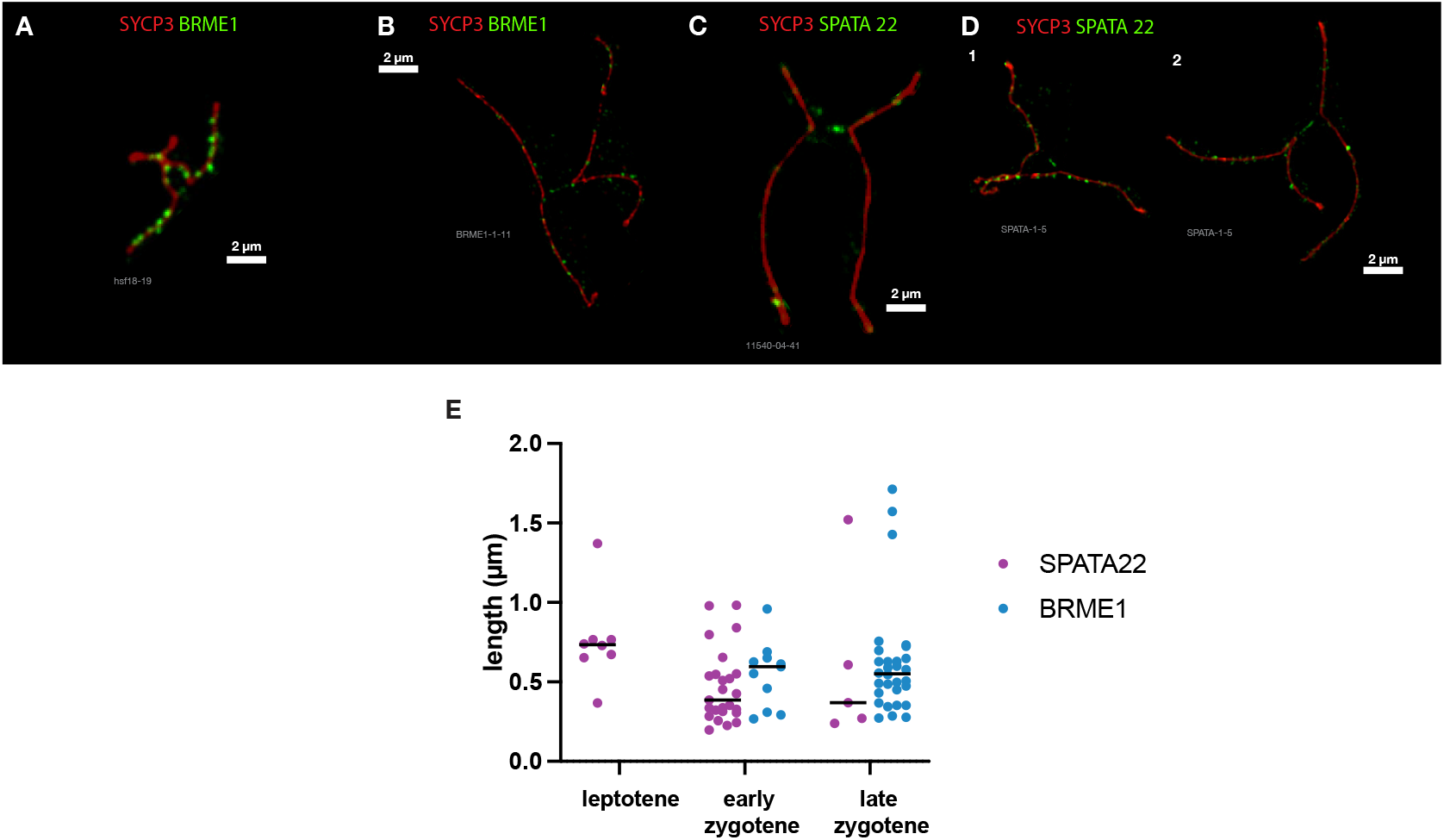
(related to Figure 5). Localization of SPATA22 and BRME1 to tensioned DNA tethers in spermatocytes. **(A)** A single example of chromosomes in butterfly arrangement connected by a tether marked with BRME1 we found in the confocal microscopy images analyzed in Figure 2I. **(B)** Examples of BRME1 on a “butterfly” structure in the SIM dataset. **(C)** Example of SPATA22-decorated tether in confocal dataset analyzed in Figure 2F. **(D)** Examples of SPATA22-decorated tether in the SIM dataset. Structure 2 is the only example of non-symmetrical butterfly we observed. **(E)** Tether lengths in SPATA22 and BRME1 SIM images. All data points and median are plotted. The experiment was performed once.

**Supplementary Figure S8.**
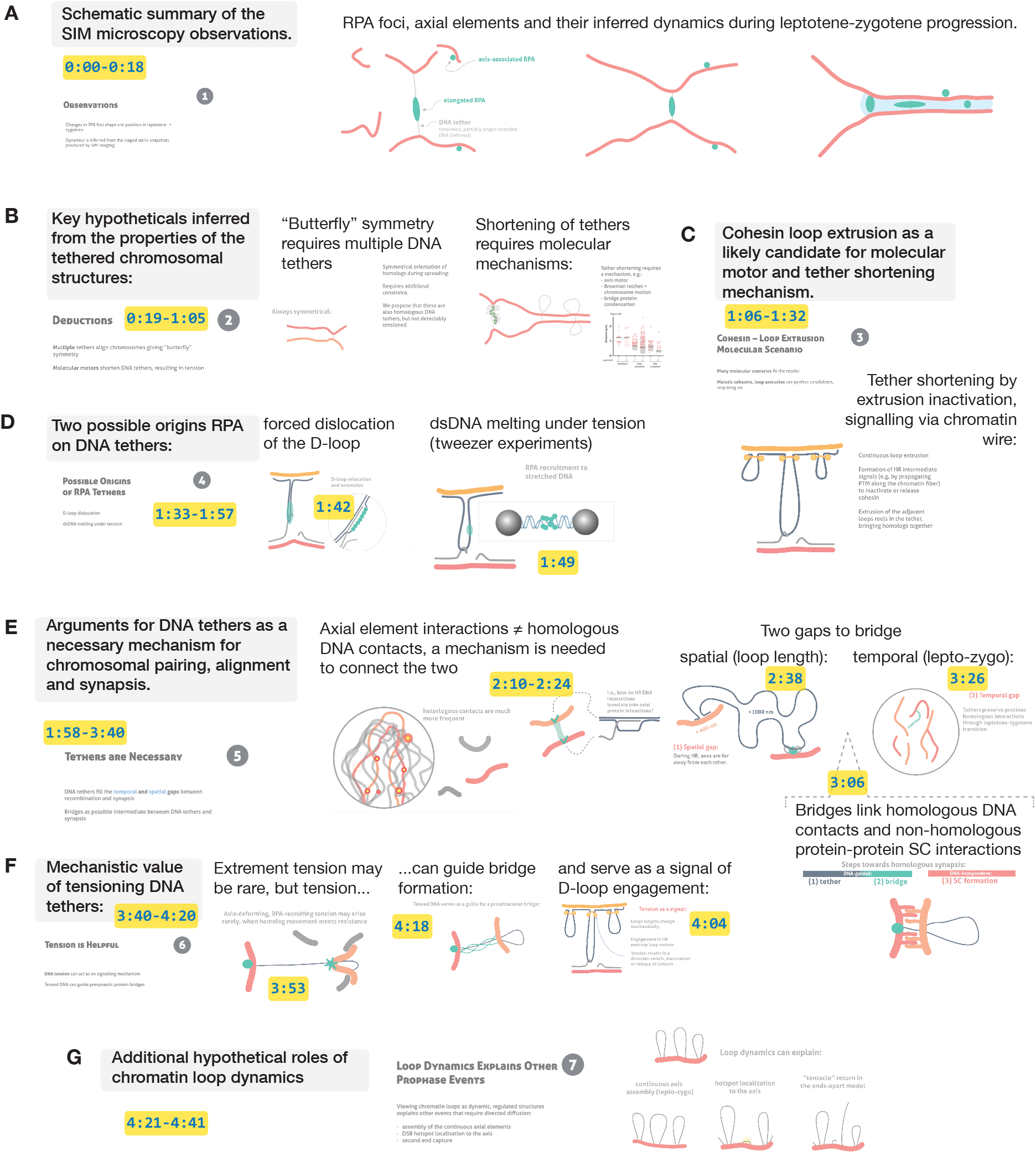
Supplementary movie annotation. Supplementary movie illustrates in a form of animation the main components of the model described in the main and the supplementary extended discussions. Panels **(A-G)** correspond to the 7 parts of the movie, with key points and screenshots of the animations. Yellow labels give references to the timestamps (min:sec).

**Supplementary Figure S9.**
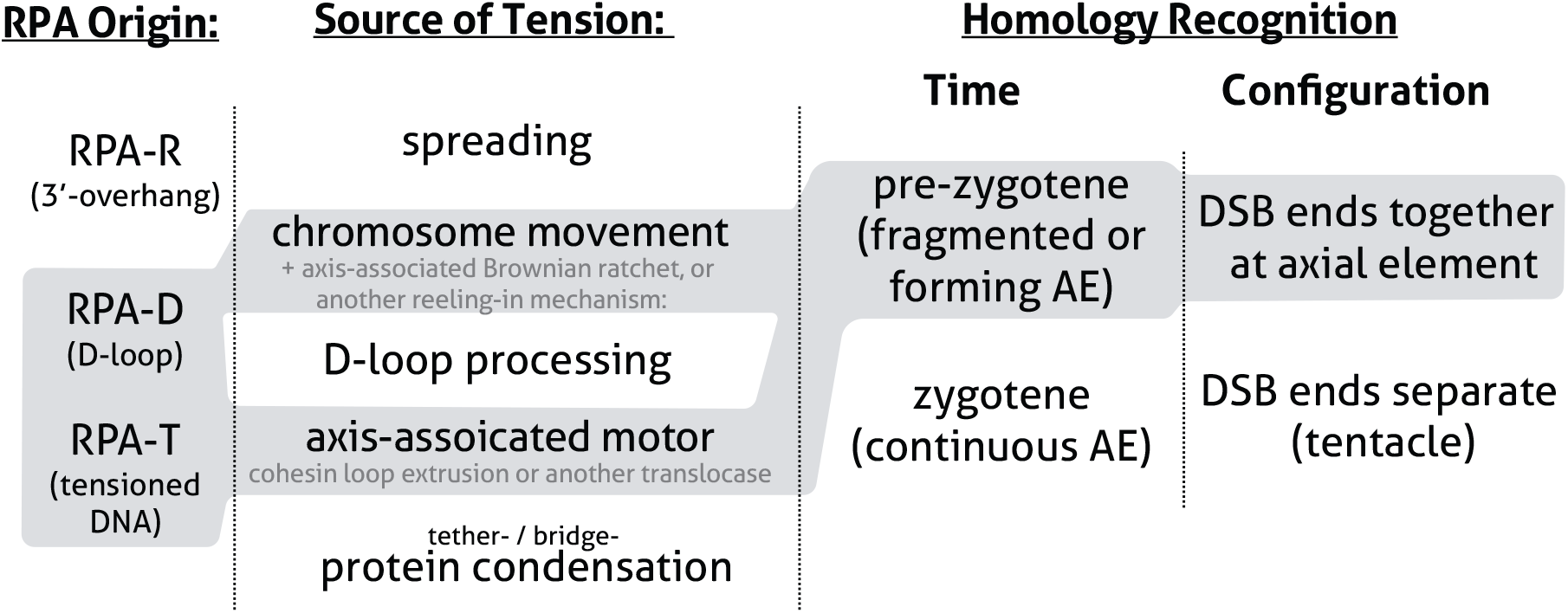
Our model is compatible with multiple molecular scenarios and their combinations. RPA may be recruited to the tether directly due to tension-induced DNA conformation change or represent a conventional HR intermediate, e.g. a D-loop within a recombinosome that dissociated from the axial element due to tension, or both. The main question is the nature of the machinery creating the tension and its localization within the tether. The most likely motor candidates are proteins that can translocate along dsDNA, anchored at the axial element. But HR intermediate remodeling (e.g. by branch migration) within the tether can also affect the length of the tether. In addition, an axis associated ratchet in combination with chromosome movement and/or protein condensation along the DNA tether are possible mechanisms for tether shortening.

